# Discovery of cancer driver genes based on nucleotide context

**DOI:** 10.1101/485292

**Authors:** Felix Dietlein, Donate Weghorn, Amaro Taylor-Weiner, André Richters, Brendan Reardon, David Liu, Eric S. Lander, Eliezer M. Van Allen, Shamil R. Sunyaev

**Affiliations:** Department of Medical Oncology, Dana-Farber Cancer Institute, Harvard Medical School, Boston, MA 02215, USA; Broad Institute of Massachusetts Institute of Technology and Harvard, Cambridge, MA, 02142, USA; Division of Genetics, Brigham and Women’s Hospital, Harvard Medical School, Boston, MA 02115, USA; Department of Biomedical Informatics, Harvard Medical School, Boston, MA 02115, USA; Koch Institute for Integrative Cancer Research, Massachusetts Institute of Technology, Cambridge, MA 02139, USA

## Abstract

Many cancer genomes contain large numbers of somatic mutations, but few of these mutations drive tumor development. Current approaches to identify cancer driver genes are largely based on mutational recurrence, i.e. they search for genes with an increased number of nonsynonymous mutations relative to the local background mutation rate. Multiple studies have noted that the sensitivity of recurrence-based methods is limited in tumors with high background mutation rates, because passenger mutations dilute their statistical power. Here, we observe that passenger mutations tend to occur in characteristic nucleotide sequence contexts, while driver mutations follow a different distribution pattern determined by the location of functionally relevant genomic positions along the protein-coding sequence. To discover new cancer genes, we searched for genes with an excess of mutations in unusual nucleotide contexts that deviate from the characteristic context around passenger mutations. By applying this statistical framework to whole-exome sequencing data from 12,004 tumors, we discovered a long tail of novel candidate cancer genes with mutation frequencies as low as 1% and functional supporting evidence. Our results show that considering both the number and the nucleotide context around mutations helps identify novel cancer driver genes, particularly in tumors with high background mutation rates.

---

Multiple algorithms have been developed to systematically identify genes that drive tumor formation^1–5^. Most search for genes harboring more nonsynonymous mutations than expected based on the local background mutation rate ^1–5^. These recurrence-based methods have successfully identified many novel cancer genes^4,6,7^. However, several studies have noted that the sensitivity of recurrence-based approaches is limited^6,8,9^, because functionally neutral passenger mutations dilute their statistical power to detect recurrent driver mutations^10,11^. Hence, due to the high prevalence of passenger mutations in tumors with high background mutation rates, recent studies have concluded that orders of magnitude more sequencing data would be needed to establish a comprehensive catalog of all cancer driver genes using recurrence-based methods^6,8,9^, (Fig. S1).

Passenger mutations are not uniformly distributed along the cancer genome. Rather, they are enriched within characteristic nucleotide sequence contexts, whose specificity depends on the specific mutational processes active in a given tumor^12–15^. For instance, APOBEC enzymes scan single-stranded DNA for specific nucleotide sequence motifs and deaminate cytidine to uracile within these motifs^16–18^. Similarly, mutant polymerase s randomly introduces passenger mutations in a non-uniform manner, since its fidelity depends strongly on the local nucleotide context^19–22^. For driver mutations, the distribution depends not only on the local nucleotide context, but also on the location of functionally relevant positions along the protein sequence^23–26^. Thus, we hypothesized that the nucleotide context would differ substantially around driver and passenger mutations.

Based on this hypothesis, we here developed a biologically informed statistical approach for discovering new cancer genes. Briefly, our approach searches for genes harboring an excess of mutations in unusual nucleotide contexts that deviate from the characteristic nucleotide context around passenger mutations (Fig. 1a, Methods).

**Fig. 1.**
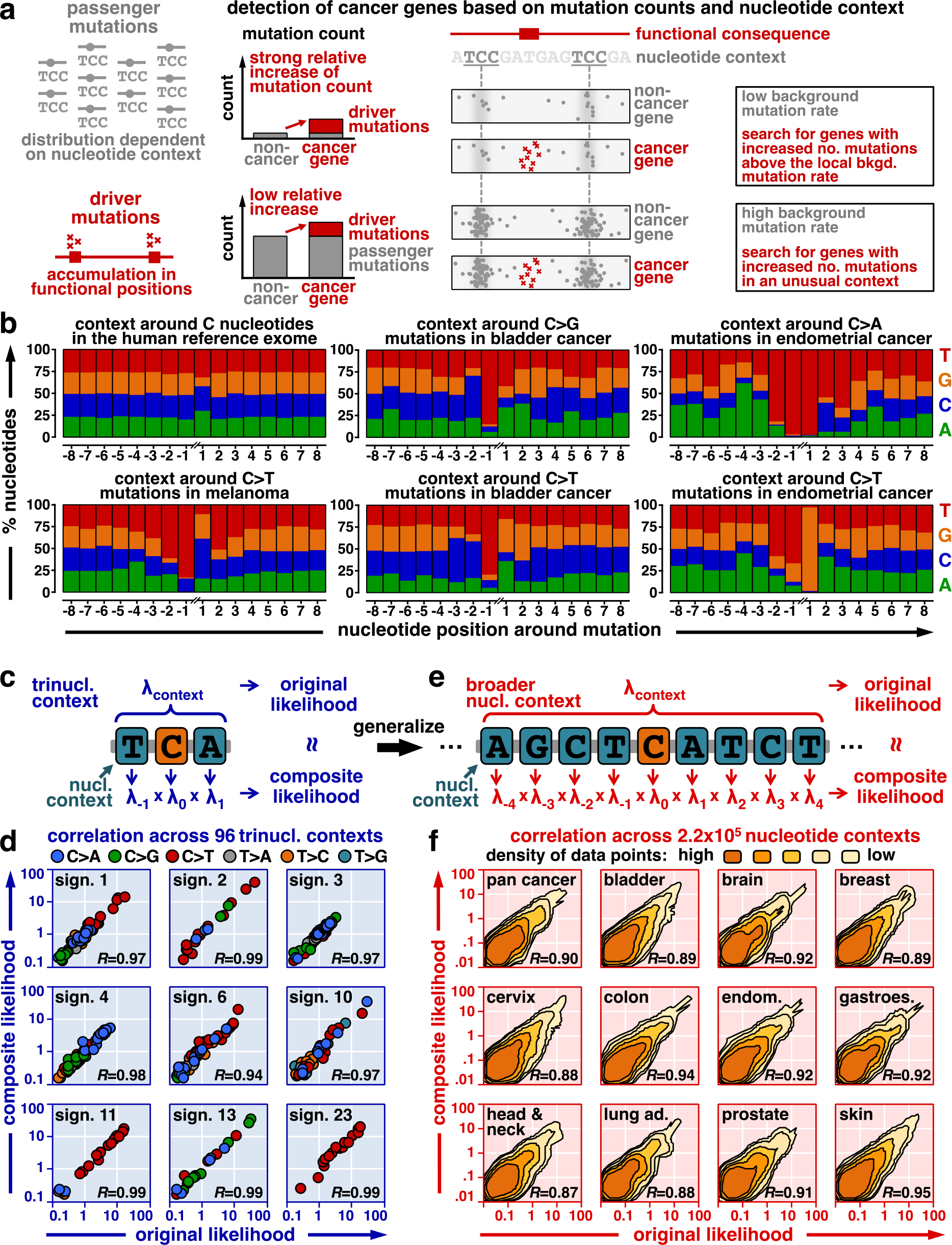
A biologically informed statistical framework to discover candidate cancer genes. **a**, Schematic of our statistical framework to discover candidate cancer genes based on nucleotide context. Passenger mutations accumulate in characteristic nucleotide contexts (gray, left), whereas driver mutations typically accumulate in functionally relevant positions (red, left). We searched for genes harboring an increased number of nonsynonymous mutations above the local background mutation rate (mutational recurrence, middle). Further, we searched for genes with an excess of mutations in nucleotide contexts that deviate from the characteristic nucleotide context around passenger mutations (mutations in unusual contexts, right). In tumors with high background mutation rates, the second criterion allowed us to actively suppress mutations in the test statistics that were likely to be passenger mutations based on their surrounding nucleotide context (gray). **b**, The nucleotide context around passenger mutations is visualized for three cancer types with high average background mutation rates. In brief, we counted how often we observed which nucleotide in the context around recurrent passenger mutations (±8 nucleotides). These plots show that the flanking 5’ and 3’ nucleotides have the strongest impact on the local mutation probability (±1, trinucleotide context). However, also flanking nucleotides outside of the trinucleotide context have a substantial impact on the local mutation probability, suggesting that the broad nucleotide context around passenger mutations contains a relevant biological signal that we needed to consider in our approach. **c-f**, To integrate this signal into our statistical framework, we developed a composite likelihood model that characterizes the broad context around passenger mutations. **c**, Mutation probabilities of trinucleotide contexts are commonly modeled by determining the mutation probability of each possible trinucleotide context independently^11–14^ (original likelihood, top). Instead, we integrated the effect of the flanking 5’ and 3’ nucleotides, as well as the base substitution type as independent factors into a composite likelihood model (bottom). **d**, For each classical trinucleotide mutation signature^11–14^, we plotted the original mutational likelihood (x-axis) against the composite likelihood (y-axis). Dot colors reflect the six different base substitution types, and Pearson correlations are annotated on the bottom right. These analyses revealed that mutation probabilities of trinucleotide contexts could be decomposed into the effects of their central and flanking 5’ and 3’ nucleotides, thus corroborating the validity of our composite likelihood approach for trinucleotide contexts. **e**, We next generalized the composite likelihood model to broader nucleotide contexts. In parallel to our approach for trinucleotide contexts, we integrated the effect of each flanking nucleotide in the broad context as an independent and multiplicative factor into the composite likelihood. **f**, We then counted the number of mutations in each possible 7-nucleotide context (x-axis, original likelihood) and compared them with the composite likelihood (y-axis). Since the number of possible nucleotide contexts was too large to be visualized directly, we plotted the data point density. Similar plots for the remaining trinucleotide signatures and cancer types are shown in Figures S7a-c. An analysis of the contribution of flanking nucleotides outside of the trinucleotide context to the local mutation probability in the composite likelihood model is shown in Figures S7d and S8.

Our new method requires modeling the nucleotide context around passenger mutations. The 5’ and 3’ nucleotides immediately adjacent to a passenger mutation have the strongest effect on the local mutation probability (Fig. 1b, S2-S3). However, as reported previously^27,28^, the additional upstream and downstream nucleotides flanking a passenger mutation also influence its mutation probability substantially (Fig. 1b, S2-S3). Traditionally, the effect of the flanking 5’ and 3’ nucleotides on the local mutation probability has been modeled by determining the mutation probabilities of all possible trinucleotide contexts independently^12–15^. As the number of flanking nucleotides increases, the number of possible sequence contexts grows exponentially - soon exceeding the number of mutations per tumor (Fig. S4). Hence, it is no longer feasible to analyze all possible sequence contexts independently.

Instead, we approximated the context-specific mutation probability by assuming that each flanking nucleotide contributed independently and multiplicatively to the local mutation probability (Fig. 1c-f, S5-S8, Methods). For instance, we approximated the mutation probabilities of trinucleotide contexts as products of the effects of their flanking 5’ and 3’ nucleotides, as well as their base substitution type (Fig. 1c-d, S7a). We developed a composite likelihood model^29^ to extend this approach to larger nucleotide contexts (Fig. 1e). This model closely matched the observed mutation probabilities for the 29 cancer types examined in this study (Fig. 1e-f, S7b-c). Although the immediately adjacent 5’ and 3’ nucleotides had the strongest impact on the local mutation probability, also flanking nucleotides outside of the trinucleotide context had a substantial effect in this composite likelihood model, thus refining our approximation of the local mutation probabilities (Fig. S7d, S8).

We then examined whether the composite likelihood model could distinguish driver from passenger mutations using 10 established melanoma genes and 5 non-cancer-related genes that had been reported as false-positive findings in previous cancer gene discovery studies^3^ (Fig. 2). While mutations in non-cancer-related genes closely followed the expected context-dependent distribution pattern derived from the composite likelihood model, most mutations in cancer genes fell in nucleotide contexts that deviated from the expectation of the model. This suggested that considering the broad nucleotide context around mutations could indeed provide new biological information to help distinguish between driver and passenger mutations.

**Fig. 2.**
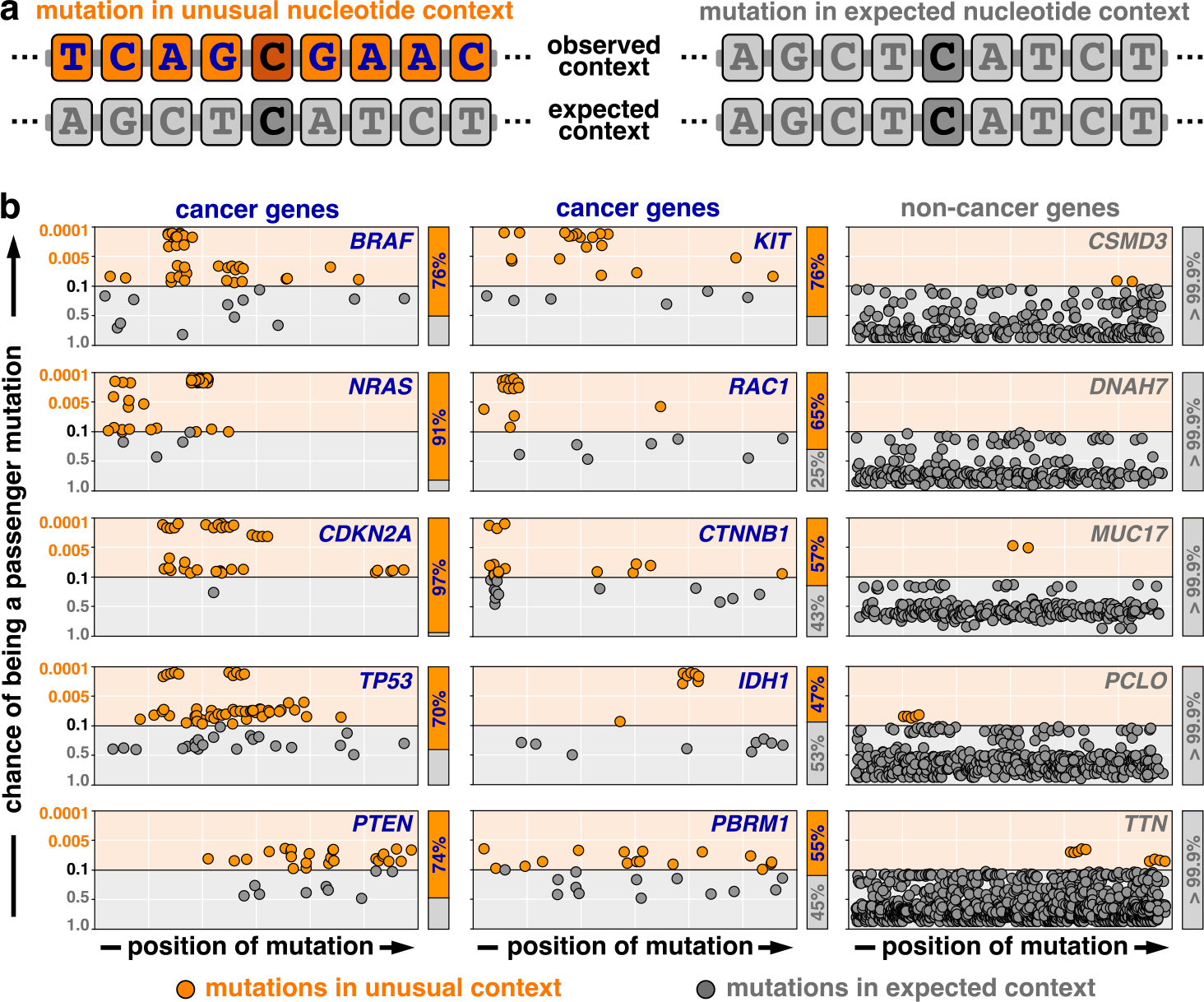
Cancer driver genes harbor an excess of mutations in unusual nucleotide contexts. **a**, For each mutation, we compared its nucleotide context (observed context, top) with the characteristic context around passenger mutations (expected context, bottom). We derived a probability score that indicated whether the mutation occurred in an unusual (left, orange) or expected (right, gray) nucleotide context (Methods). **b**, We corrected these probabilities for multiple hypothesis testing (false-discovery rates, y-axis) and plotted them against their genomic position (x-axis). In cancer genes a substantial number of mutations occurred in unusual sequence contexts (left, middle). In non-cancer genes mutations in unusual sequence contexts were extremely rare (right). This suggested that cancer driver genes harbor an increased number of mutations in unusual nucleotide contexts that deviate from the characteristic nucleotide context around passenger mutations. This observation provides a novel biological criterion to discriminate between driver and passenger mutations.

Encouraged by these observations, we developed a statistical framework to detect cancer driver genes that considers both mutation counts and nucleotide contexts. In our model, the probability of observing the number *n*_*g*_ and the context-dependent distribution *ν*_*g*_ of nonsynonymous mutations in a gene *g* (*P*(*n*_*g*_,*ν*_*g*_|*s*_*g*_;*λ*_*g*_)) depends on the number of synonymous mutations *s*_*g*_ and the context-specific mutation rates *λ*_*g*_. We decomposed this probability into the probability of observing *n*_*g*_ nonsynonymous mutations, given the number of synonymous mutations *s*_*g*_ (“mutation count”; *P*(*n*_*g*_|*s*_*g*_)), and the probability of these *n*_*g*_ nonsynonymous mutations falling in nucleotide contexts *ν*_*g*_, given their context-specific mutation rates *λ*_*g*_ (“nucleotide context”; *P*(*ν*_*g*_|*n*_*g*_;*λ*_*g*_)):

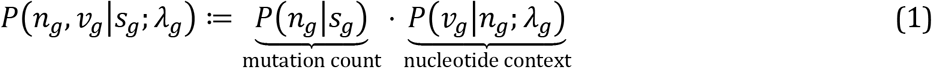

Here, *P*(*n*_*g*_|*s*_*g*_) reflects the established statistics used by existing recurrence-based methods for cancer gene discovery^1–5^. The *p*-value of *P*(*ν*_*g*_|*n*_*g*_; *λ*_*g*_) was derived by comparing the observed nucleotide contexts *ν*_*g*_ against a large number of random scenarios generated by a Monte Carlo simulation approach based on the same the context-specific mutation rates *λ*_*g*_ ^30,31^. As shown by Q-Q-plots^32^, the *p*-values derived from *P*(*n*_*g*_|*s*_*g*_), *P*(*ν*_*g*_|*n*_*g*_; *λ*_*g*_), and *P*(*ν*_*g*_,*n*_*g*_|*s*_*g*_; *λ*_*g*_) closely approximated a uniform distribution, which indicated that our models were reasonably well calibrated to the observed data (Fig. 3a, Methods).

**Fig. 3.**
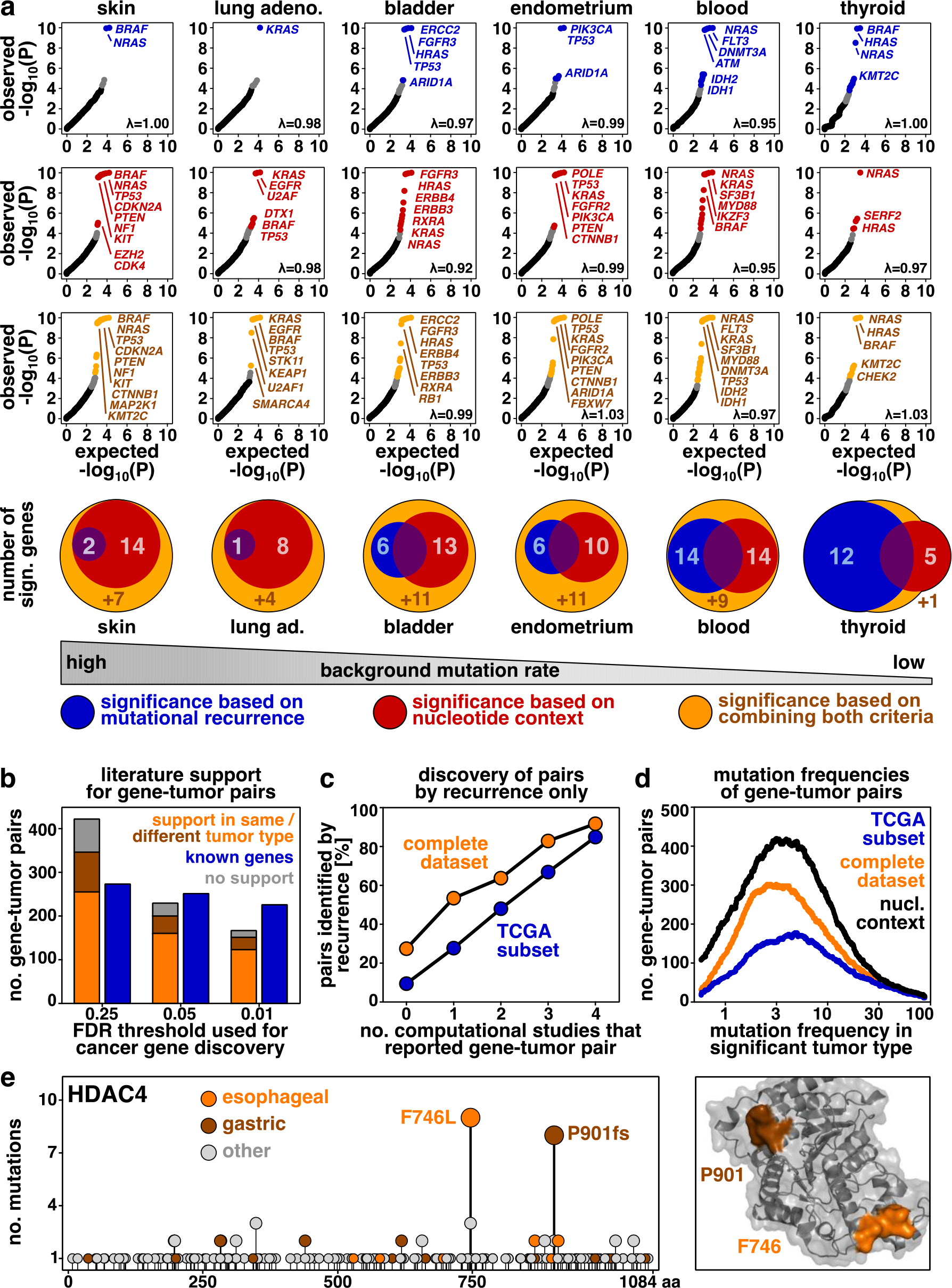
Discovery and characterization of candidate cancer genes identified based on nucleotide context. **a**, We determined which genes emerged as significantly mutated (false-discovery rate, FDR<0.25) based on their mutational recurrence (blue) and based on their excess of mutations in “unusual” nucleotide contexts (red). Further, we identified candidate cancer genes based on a statistical model that combined mutational recurrence and nucleotide context (orange). We compared the expected (x-axis) and observed (y-axis) *p*-values derived from these three statistical models using Q-Q-plots. Venn diagrams visualize the overlap in significant genes detected with these three models (bottom). These analyses revealed that increased mutation counts and unusual nucleotide contexts provide two complementary criteria for the discovery of cancer genes. Integrating both aspects into a combined significance model enabled discovery of candidate cancer genes across tumor types with high and low background mutation rates (left to right). **b**, We stratified our findings based on their support in the literature. Known gene-tumor pairs, which had been reported as significantly mutated previously, are colored in blue. Novel gene-tumor pairs, which had not been reported as significantly mutated previously, are colored in orange (experimental support in the same tumor type), brown (literature support in a different tumor type), or gray (no support). For rigorous FDR thresholds (FDR<0.01), a majority of the significant gene-tumor pairs (82%, 323/395) involved canonical cancer genes in the Cancer Gene Census^34’35^. Further, most gene-tumor pairs had been known previously or had experimental literature support in the same tumor type (89%, 351/395 for FDR<0.01). For less stringent FDR thresholds, the absolute number of novel findings with experimental literature support increased, and the number of findings without literature support (11%, 75/697) did not exceed the expected false-discovery rate (FDR<0.25). **c**, We counted for each gene-tumor pair (FDR<0.25) how many previous studies reported the gene-tumor pair as significantly mutated^4,6,7,33^ (x-axis). Further, we examined whether the gene-tumor pair was also identified using an established recurrence-based approach^3^ (y-axis). The concordance between these two measures potentially reflects the fact that most previous pan-cancer gene discovery studies used recurrence-based approaches to identify cancer genes^4,6,7,33^. **d**, We explored the mutation frequencies of the gene-tumor pairs that emerged as significantly mutated based on their recurrence in the TCGA subset (blue), in the complete dataset (orange), or when additionally considering the nucleotide context around mutations (black). This density plot revealed that both the addition of 4,913 samples from TCGA-independent studies and considering the nucleotide context around mutations independently contributed to the discovery of rare candidate cancer genes. e, Exemplary evidence for the candidate cancer gene *HDAC4*. Left: The distribution of *HDAC4* mutations is visualized as a needle plot. For each amino acid substitution the number of samples (y-axis) is plotted against its position in the peptide sequence (x-axis). Dot colors reflect the tumor types, in which the amino acid substitution was detected. Right: The position of the two mutational hotspots is visualized using a crystal structure^42^ (PDB: 4CBY). A previous study reported a hydrogen bond and salt bridge network between W762, E764, and R730, which along with F746 form a closed hydrophobic patch peripheral to the catalytic center of HDAC4^43^ (orange). Evidence for other candidate cancer genes can be found in Figures S11-S13.

Notably, mutational count and nucleotide context provided complementary criteria for detecting cancer genes (Fig. 3a). In cancer types with low background mutation rates, such as thyroid cancer, mutational counts were highly informative. In cancer types with high background mutation rates, such as melanoma, the nucleotide context was the dominant criterion. Combining both criteria identified several candidate cancer genes that could not be identified based on mutational count or nucleotide context alone (Fig. 3a).

We applied our statistical framework to whole-exome sequencing data from 12,004 individual tumors spanning 29 different tumor types (Fig. S9, Table S1). The results of these analyses are summarized here (Fig. 3-4, S9-45) and at www.cancer-genes.org, for various false-discovery rate (FDR) thresholds. For FDR<0.25, we identified 697 gene-tumor pairs, i.e. pairs of significantly mutated genes and their associated tumor type. These gene-tumor pairs involved 379 distinct genes, with 423 gene-tumor pairs being novel. The corresponding numbers were 484, 252 and 231 for FDR<0.05, as well as 395, 201, and 168 for FDR<0.01 (Tables 1, S2-S3). Gene-tumors pairs were considered novel if they were not reported as significantly mutated in at least two computational studies, among all TCGA marker papers, a meta-analysis of 876 publications, and two large-scale pan-cancer gene discovery studies^4,6,7,33^.

**Table 1.** Stratification of candidate cancer genes by literature support. To examine the biological relevance of our findings, we stratified them based on their literature support. Genes and gene-tumor (G-T) pairs that had been reported as significantly mutated in at least two previous computational studies were classified as known (blue, 2nd row). Novel genes and gene-tumor pairs, which had not been reported as significantly mutated previously (red, 3rd row), were further stratified depending on whether there was literature support (experimental or clinical) for the same tumor type in which we discovered them as significantly mutated (orange, 4th row), supporting literature for a different tumor type (brown, 5th row), or no supporting data (gray, 6th row). Depending on their literature support level, 94% (known, 257/274), 72% (same tumor type, 186/257), 23% (different tumor type, 21/91), and 3% (no support, 2/75) of the gene-tumor pairs (FDR<0.25) involved canonical cancer genes present in the Cancer Gene Census^34,35^, compared with a rate of 3.8% for random gene-tumor pairs. Thus, literature support levels provide a measure to prioritize our findings based on their external validity.

|  | FDR < 25% |  | FDR < 10% |  | FDR < 5% |  | FDR < 1% |  |
| --- | --- | --- | --- | --- | --- | --- | --- | --- |
|  | Genes | G-T Pairs | Genes | G-T Pairs | Genes | G-T Pairs | Genes | G-T Pairs |
| **Total** | **379** | **697** | **298** | **562** | **252** | **484** | **201** | **395** |
| **Known** | **127** | **274** | **123** | **262** | **121** | **253** | **114** | **227** |
| **Novel** | **252** | **423** | **175** | **300** | **131** | **231** | **87** | **168** |
| ● support in same type | 110 | 257 | 85 | 198 | 70 | 161 | 50 | 124 |
| ● support in different type | 69 | 91 | 44 | 55 | 31 | 40 | 21 | 28 |
| ● no support | 73 | 75 | 46 | 47 | 30 | 30 | 16 | 16 |

We next examined the biological relevance of the 423 novel gene- tumor pairs (FDR<0.25). Half of the novel gene-tumor pairs (49%) involved canonical cancer genes in the Cancer Gene Census^34,35^, compared with a rate of 3.8% for random gene-tumor pairs. We systematically reviewed the literature to further investigate the experimental or clinical support for of the novel gene-tumor pairs. We only considered publications with experimental data supporting the causal involvement of these genes in carcinogenesis and excluded functionally unsupported reports of mutations (Methods). A majority of the novel gene-tumor pairs (82%) had experimental support, with 61% having support in the same tumor type, in which we detected them as significantly mutated (Fig. 3b, Tables 1, S2-S3). In contrast, the rate for random gene-tumor pairs was 17%. Overall, 11% (75/697, FDR<0.25), 6% (30/484, FDR<0.05), and 4% (16/395, FDR<0.01) of the significant gene-tumor pairs had no literature support, which is roughly in accordance with these FDR thresholds (Fig. 3b, Tables 1, S2-S3).

We asked whether considering the nucleotide context identified candidate cancer genes that were not discovered based on recurrence alone. Among gene-tumor pairs previously reported as significantly mutated, 74% were also identified by using an established recurrence-based approach^3^ (FDR<0.25 for both methods, Fig. 3c-d, S10). In contrast, among novel gene-tumor pairs, only 33% were identified based on recurrence alone. In particular, our statistical framework identified numerous biologically relevant candidate cancer genes that were not identified based on recurrence alone. For instance, *HDAC4* (histone deacetylase 4) was significantly mutated in gastroesophageal cancer (FDR=5.5×10^−2^ by nucleotide context and recurrence; FDR=6.8×10^−1^ by recurrence alone; not reported as significant previously; Fig. 3e, S11). Histone deacetylases have been implicated in tumor formation^36^–^38^ and *HDAC4* displayed two mutational hotspots: gastric cancers with disruptive frameshift mutations (P901fs), and esophageal cancers with recurrent missense mutations (F746L) (Fig. 3e). Similarly, we identified *POLR2A* (RNA polymerase II subunit A) as significantly mutated in lung adenocarcinoma (FDR=1.07×10^‒5^ by nucleotide context and recurrence; FDR=1.0 by recurrence; not reported as significant previously; Fig. 4, S12). Mutations in *POLR2A* have been implicated in the development of meningioma^39^, and *POLR2A* has been identified as a therapeutic target in colon cancer due to its frequent co-deletion with *TP53*^40^. Further, we noticed that *POLR2A* contained recurrent mutations in positions that are relevant for the protein-DNA interaction (Fig. S12). Additional biologically relevant candidate cancer genes that were not identified based on recurrence included *ANAPC1*, *FGFR4*, *IKZF3*, *PARG*, *SOX17*, and *ZFX* (FDR<0.1 by nucleotide context and recurrence; FDR=1.0 by recurrence; Fig. 4, S11-S12, Tables S2-S3). In addition, we observed that the following cancer-related signaling complexes contained several candidate cancer genes, i.e. new cancer genes or gene-tumor pairs: modulation of Ras signaling (*RHOA*, *RHOB*, *RRAS2*), cell cycle regulation (*CCNQ*, *CDK4*), regulation of protein levels (*EEF1A1*, *EIF1AX*, *MIA2*), the catenin/cadherin complex (*FAT1*, *FAT3*, *FAT4*), DNA polymerases (*POLQ*, *POLR2A*, *REV3L*), regulation of transcription (*MAML2*, *SF3B2*), modulation of apoptosis (*ACVR2A*, *ACVR1B*, *CASP8*, *BIRC3*, *BIRC6*), and epigenetic modification (Fig. S13). In these signaling complexes, 64% (118/183) of the gene-tumor pairs had not been reported as significantly mutated previously, and 60% (110/183) of the gene-tumor pairs were not identified by recurrence alone (Fig. 4).

**Fig. 4.**
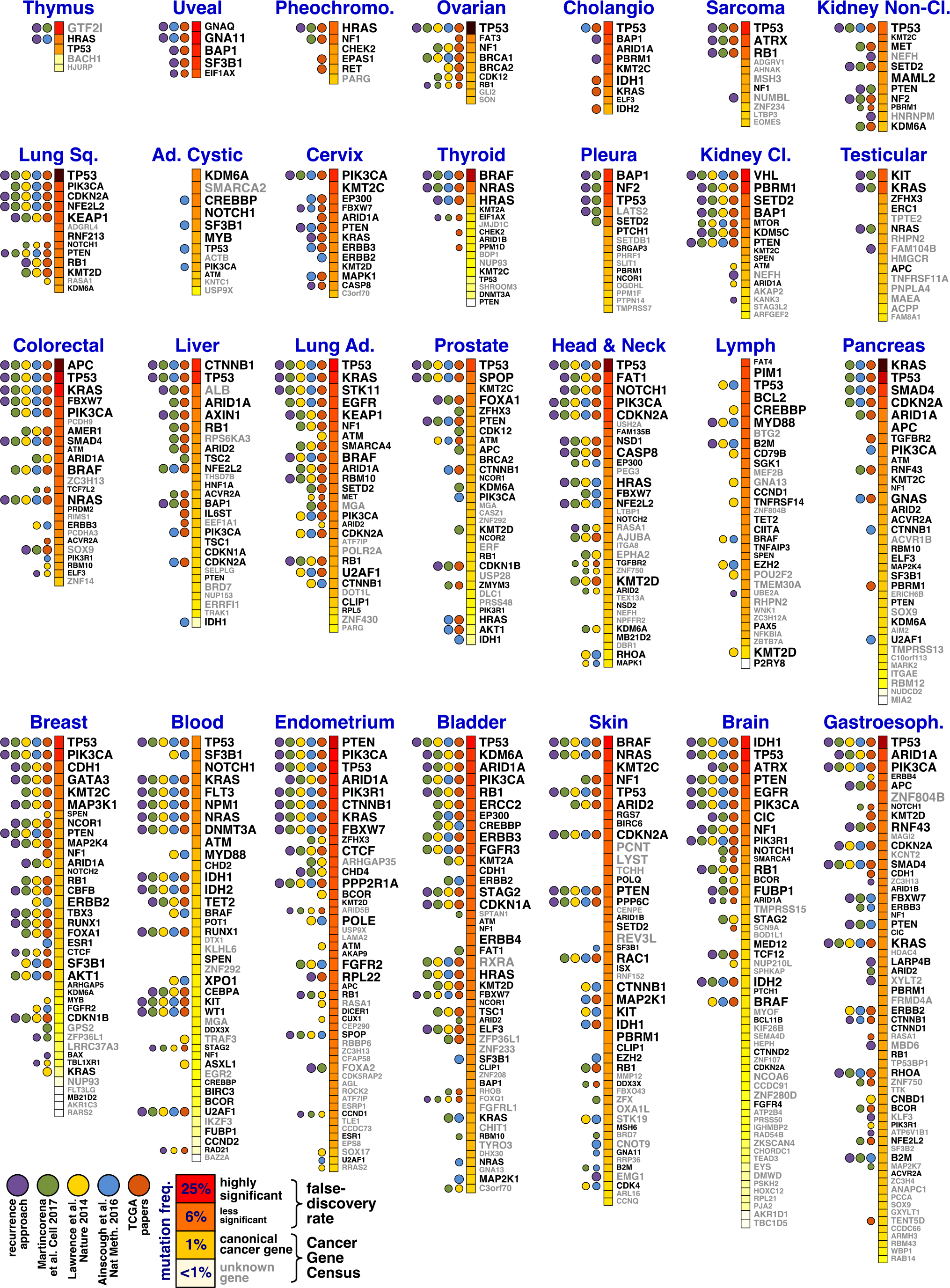
A refined catalog of driver genes involved in human cancer. We applied our statistical framework to whole-exome sequencing data from 12,004 tumors. Significant gene-tumor pairs (FDR<0.25) are listed in decreasing order according to their mutation frequency, which is reflected by the color of the square next to the gene name (dark red to white). The font size of the gene name reflects its significance (false-discovery rate), and the font color (black vs. white) indicates whether the gene is a canonical cancer gene in the Cancer Gene Census^34,35^. To determine which gene-tumor pairs had been known previously, we benchmarked our results against all TCGA marker papers^7^ (orange), a meta-analysis^33^ of 876 publications (blue), the tumorportal database^6^ (yellow), and a pan-cancer study, which adopted the dN/dS ratio for cancer gene discovery^4^ (green). We further ran an established recurrence-based approach^3^ on our dataset (purple) to determine which gene-tumor pairs were identified based on recurrence alone. A more detailed overview of the driver mutation landscape of individual tumor types is provided in Figures S18-S45. An interactive visualization of these results can be found online (www.cancer-genes.org).

Taken together, our findings demonstrate that characterization of the broad nucleotide context around somatic passenger mutations enhances cancer gene discovery, particularly in tumor types with high background mutation rates. Consideration of the nucleotide context for cancer gene discovery does not require prior knowledge of the location of functionally relevant positions or the biological effect of mutations. Hence, nucleotide contexts may ultimately be amenable to variant and gene discovery in non-coding regions of the cancer genome. Through our statistical model we identified a long tail of reasonable candidate cancer genes that may form the foundation for future experimental and clinical studies. The new statistical framework is available as a fully executable software tool called MutPanning (www.cancer-genes.org) and can be executed online as a module on the GenePattern platform^41^ (www.genepattern.org).

## Supporting information

## Acknowledgments

We thank Dr. Gad Getz for valuable comments and suggestions. F.D. was supported by EMBO (ALTF 502-2016). E.M.V.A. and S.R.S received funding from the National Institutes of Health (K08 CA188615, R01 CA227388 to E.M.V.A; R01 MH101244, R35 GM127131, U01 HG009088 to S.R.S.). E.M.V.A further acknowledges support by the Phillip A. Sharp Innovation in Collaboration Award.

