## Supplementary material for "Discovery of cancer driver genes based on nucleotide context"

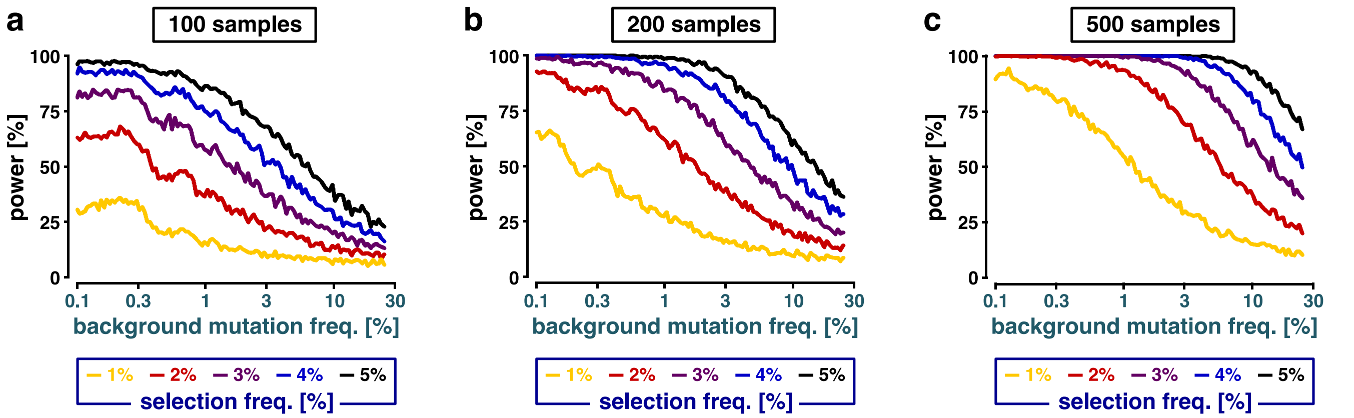


**Extended Data Fig. S1 | Passenger mutations limit the ability of recurrence-based approaches to detect rare cancer driver genes. a-c,** These analyses support the schematic shown in Figure 1a. To understand the effect of passenger mutations on the statistical power of recurrence-based approaches, we used the following toy model. Given $N$ samples and a background mutation frequency $f_{\text{bkgd}}$, we modeled the total number of passenger mutations in a non-cancer-related gene using a Poisson distribution, $\text{Pois}(N \cdot f_{\text{bkgd}})$. Further, given a positive selection frequency $f_{\text{sel}}$ (i.e., the mutation frequency of a cancer driver gene above the background mutation rate) we assumed that the number of driver mutations followed a Binomial distribution, $\text{Binom}(N, f_{\text{sel}})$. Based on these two distributions, we simulated the number of mutations in cancer-related (driver mutations + passenger mutations) and in non-cancer related genes (passenger mutations only). We then determined the statistical power to discover a cancer gene as the fraction of simulation experiments in which the simulated mutation counts in cancer- and non-cancer-related genes differed significantly. We performed these simulation experiments with various background mutation frequencies $f_{\text{bkgd}}$ (x-axis), for different selection frequencies above background $f_{\text{sel}}$ (colors of the curves), and for different cohort sizes $N$ (**a,** $N$=100 samples; **b,** $N$=200 samples; **c,** $N$=500 samples). When the background mutation frequency was smaller than the positive selection frequency, the statistical power was relatively stable. However, as soon as the background mutation frequency exceeded the positive selection frequency, the statistical power dropped rapidly. This observation is in concordance with previous studies on this issue and might explain why the detection of rare cancer genes with mutation frequencies <5% was challenging in previous studies using recurrence-based approaches. Based on this observation, we concluded that additional biological criteria would be needed to systematically discover and characterize rare cancer genes involved in tumor development.

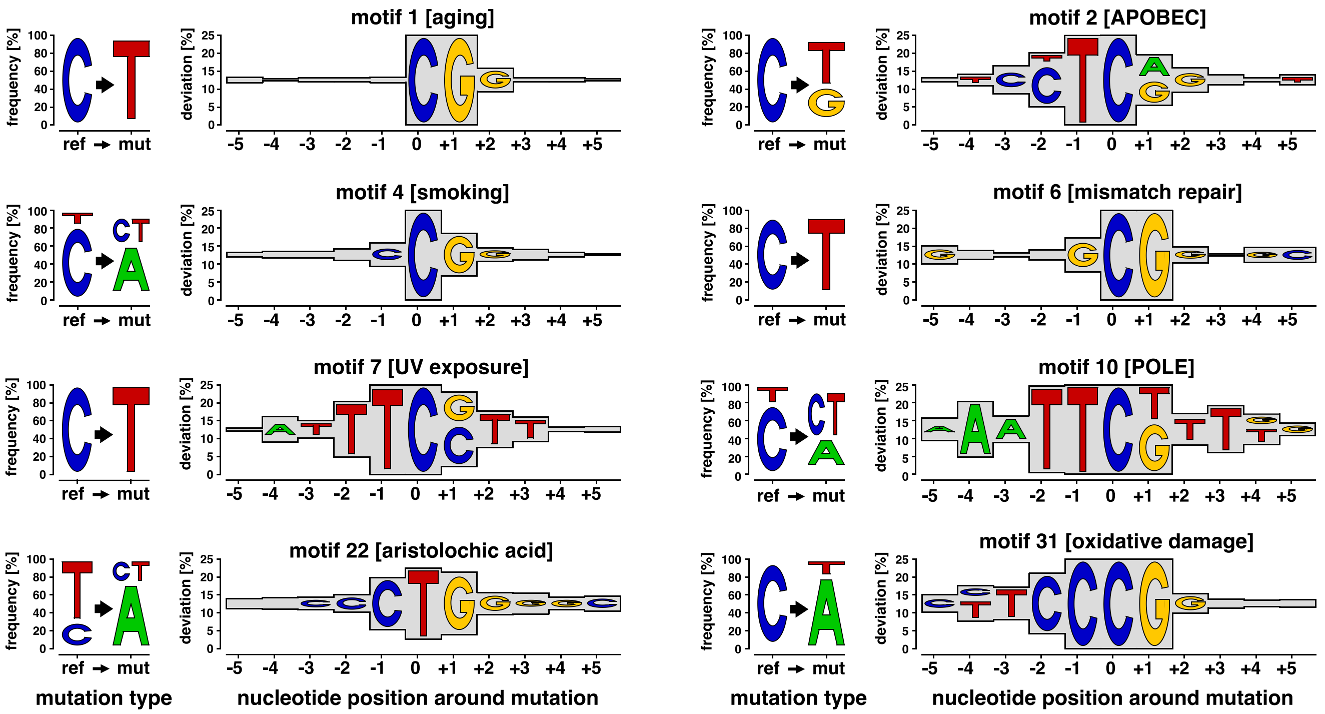


**Extended Data Fig. S2 | Passenger mutations are enriched in specific nucleotide contexts.** We observed that passenger mutations typically occured in characteristic nucleotide contexts. We visualized these characteristic contexts by sequence logo plots. Left: For each mutational process, mutation frequencies of the reference (ref) nucleotides (C vs. T) and mutation () types (mut : transitions (C/T), type-I-transitions (A), type-II-transitions (G)) are visualized as logo plots. Right: Sequence logo plots represent the relative over-representation (deviation) of the nucleotides around mutations relative to their expected frequencies in the human exome. In other words, the height of each flanking nucleotide indicates its impact on the local mutation probability. Most characteristic nucleotide contexts clearly exceed the trinucleotide context (±1) around mutations. For instance, the nucleotide context associated with UV exposure (motif 7) contains almost exclusively C>T mutations; T's are overrepresented in 5' adjacency and C/G followed by T's are overrepresented in 3' adjacency. Hence, sequence logo plots provide a convenient way to visualize the dependence of local mutation probabilities on the broad nucleotide context.

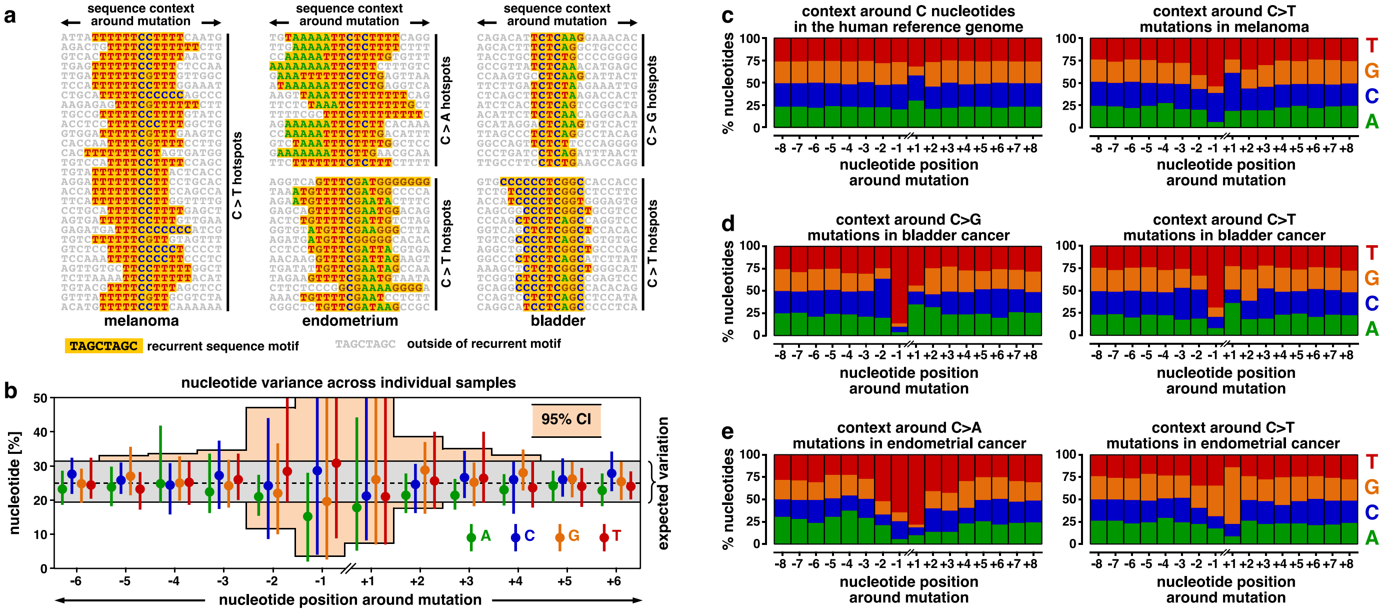


**Extended Data Fig. S3 | Initial exploration of the nucleotide context around passenger mutations.** Context-specific mutation rates are typically modeled based on the 5' and 3' nucleotides immediately adjacent to the mutation, i.e. the trinucleotide context. We explored the nucleotide context around passenger mutations and examined whether also flanking nucleotides beyond the trinucleotide context had a substantial impact on the local mutation probability. **a,** We observed that passenger mutations were enriched in characteristic nucleotide contexts, particularly for melanoma, endometrial and bladder cancer. Exemplary reads around mutations that contained these characteristic contexts (i.e. recurrent sequence motifs, yellow) are shown. This suggested that there was a substantial biological signal in the broad nucleotide context beyond the trinucleotide context. **b,** To quantify the biological signal in the broad nucleotide context, we examined the variation of the nucleotide context around mutations in samples carrying at least 500 mutations. We counted for each sample how often we observed which nucleotide in the context of its mutations (green: A, blue: C, orange: G, red: T). Based on these counts, we derived the variation of the nucleotide contexts around mutations between samples. Dots indicate the medians of these distributions, and vertical bars represent their 5$^{th}$-95$^{th}$ percentile ranges. We then examined whether this variation exceeded our expectation, assuming there was not any biological signal in the nucleotide context (beta distribution around 25% with at least 500 mutations per sample, gray envelope). The nucleotide variation (95% confidence interval over all nucleotides, orange envelope) significantly exceeded the expected variation within a ±4-nucleotide window. This suggests that there is a relevant biological signal in the flanking nucleotides outside of the trinucleotide context. **c-e,** These plots supplement Figure 1b. The nucleotide context around passenger mutations is visualized for three cancer types with high average background mutation rates. In brief, we counted how often we observed each nucleotide around non-recurrent mutations (±8 nucleotides). Based on these counts, we then determined the relative nucleotide frequencies in the nucleotide context around passenger mutations (y-axis). These plots suggest that the trend that we observed in Figure 1b also continued for non-recurrent mutations.

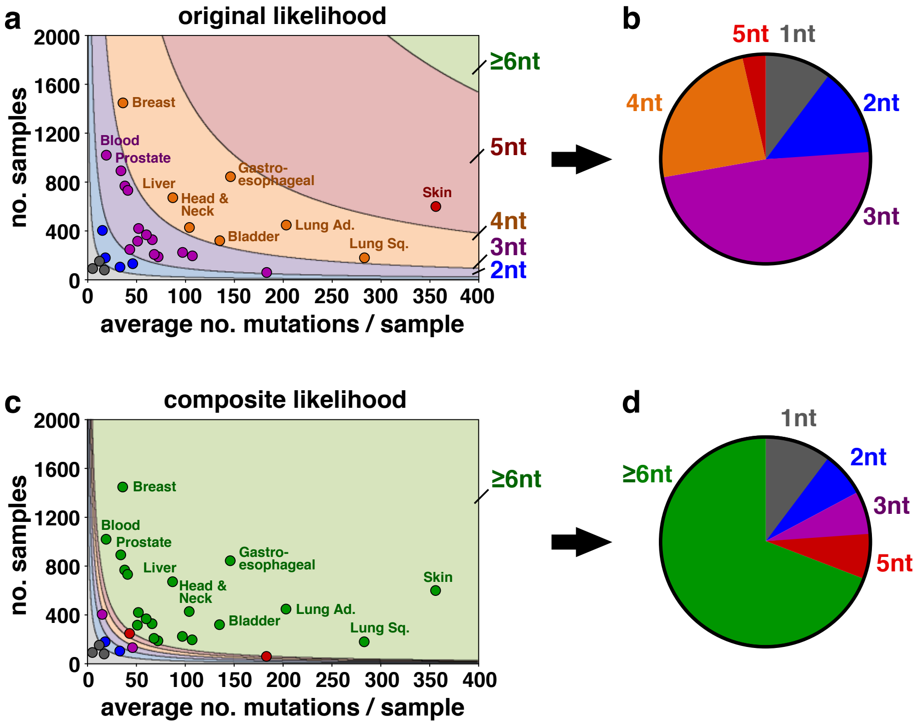


**Extended Data Fig. S4 | A new mathematical model is needed to characterize the broad nucleotide context around passenger mutations.**
**a-b,** Trinucleotide contexts (i.e., the 5' and 3' nucleotides immediately adjacent around a mutation) are commonly used to model context-dependent mutation probabilities. For this purpose, the mutation probabilities of all possible trinucleotide contexts are determined independently. However, as the number of flanking nucleotides increases, the number of possible nucleotide contexts grows exponentially. For instance, there are 96 possible trinucleotide contexts, but 24,576 possible 7-nucleotide contexts. We assumed that a fixed number of observations (arbitrarily set to 100 mutations per degree of freedom in this figure) are required to characterize each variable sufficiently. As the number of possible contexts soon exceeds the number of mutations per tumor, the traditional approach (characterizing each possible context independently, original likelihood) is intrinsically limited to the trinucleotide contexts for most cancer types. **c-d,** The composite likelihood model integrates the effect each flanking nucleotide in the context around passenger mutations as an independent factor. Hence, the degrees of freedom increase linearly with the number of flanking nucleotides included into the composite likelihood model. This allowed us to characterize a context of up to 20 nucleotides around passenger mutations. **a, c,** For each cancer type, the average number mutations per sample (x-axis) is plotted against the number of samples included in our study cohort (y-axis). Dot colors indicate the number of nucleotides, which can be integrated using the original likelihood (**a**) or the composite likelihood (**c**), respectively. **b**, **d,** Pie charts show the number of cancer types grouped according to the length of the mutation context in nucleotides, using the original likelihood (**b**) or the composite likelihood (**d**), respectively.

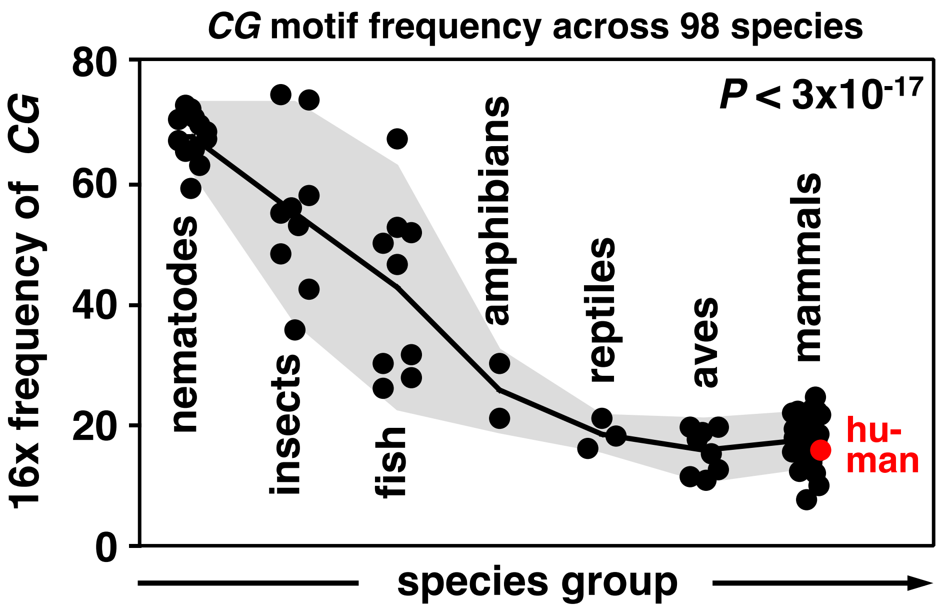


**Extended Data Fig. S5 | *CG* suppression in the human exome.** We noticed that *CG* dinucleotides were markedly underrepresented in the human exome, compared with the other 15 possible dinucleotides (red). We hence set out to examine the relative frequency of the *CG* dinucleotide across 98 different species from 7 different species groups (x-asis). Each black dot represents the relative frequency of the *CG* motif in an individual species, multiplied by 16 to normalize against the uniform distribution (y-axis). The black line connects the species group averages, and the gray envelope depicts the standard deviation for each group. These analyses suggested that the human exome displays a marked *CG* suppression, gradually evolving over the evolution. This *CG* suppression needs to be considered in our composite likelihood model.

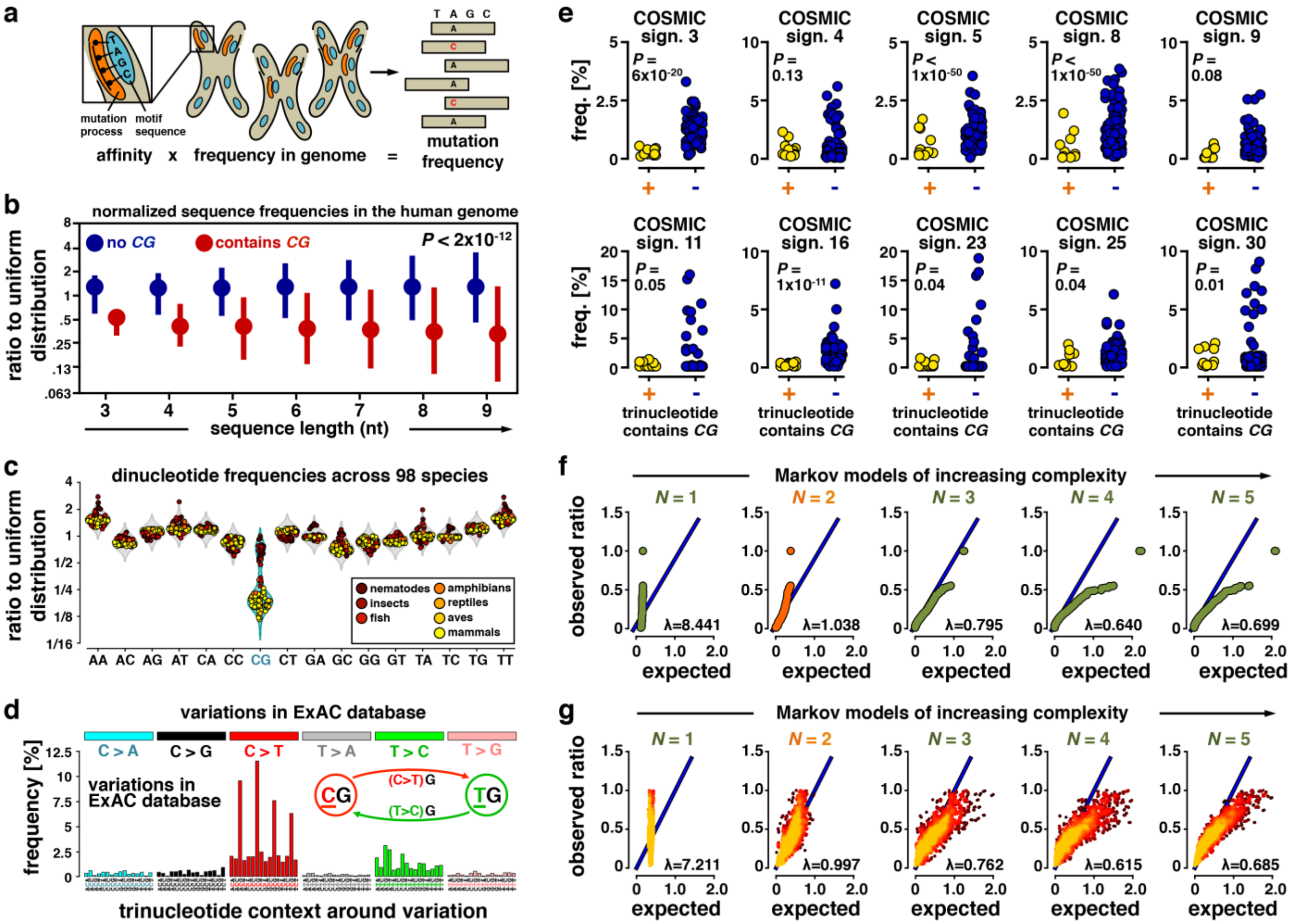


**Extended Data Fig. S6 | Characterization and modeling of the nucleotide sequence composition of the human exome.** A major prerequisite for the development of the composite likelihood model was an accurate characterization of the nucleotide sequence composition of the human exome. In other words, we aimed to develop a model that determined how often each possible nucleotide sequence occurred in the human exome. In **a-e** we describe that the sequence composition of the human exome is non-trivial, i.e. that different sequences occur at different frequencies. For instance, the human exome is characterized by a marked suppression of *CG* dinucleotides. In **f-g** we develop a Markov model to model the sequence composition of the human exome.

**a,** Schematic depicting the relationship between mutational likelihood and the human exome. For each possible nucleotide sequence, the composite likelihood model describes whether it is over- or underrepresented around passenger mutations compared with its occurrence in the human exome. In other words, for each possible nucleotide context, the composite likelihood delivers a frequency ratio, where a ratio > 1 indicates that the nucleotide context has a higher frequency around passenger mutations compared with its frequency in the human exome (the “affinity” of the mutational process to a nucleotide sequence). **b,** *CG* dinucleotide motifs are underrepresented in the human exome. We determined the relative frequency of each possible nucleotide sequence (x-axis: 3 to 9 nucleotides, nt) in the human exome. We normalized the frequencies to the total number of sequences of the same length (y-axis: x-fold change relative to a uniform distribution). For each sequence length, we plotted the distribution of the ratios: medians are represented by dots, and vertical lines plot the 5-95% quantiles. Ratios fluctuated around 1 for motifs without *CG* (blue), irrespective of the sequence length. Nucleotide sequences that contained at least one *CG* dinucleotide (red) were significantly and substantially underrepresented. Hence, nucleotide sequences are not equally represented in the human exome and the *CG* underrepresentation had to be considered in the composite likelihood model. **c,** We determined the relative representation of the 16 dinucleotide motifs (y-axis) in the genomes of 98 species from 7 different species groups (x-axis). Frequencies were normalized to the total number of dinucleotide motifs, so that a frequency ratio of 1 reflected the uniform distribution. Violin plots show that there was a bimodal distribution pattern for the *CG* dinucleotide motif across species, whereas the frequency ratios closely varied around 1 for the other dinucleotide motifs. The bimodal distribution for *CG* was clearly separated between different species groups, suggesting that this motif was lost during evolution. **d,** Based on the Exome Aggregation Consortium (ExAC) project, which contains whole-exome sequencing data from 60,706 unrelated individuals, we characterized the nucleotide context around common single nucleotide variations from the reference genome. We determined the trinucleotide context and substitution type for each of these common variants. Their genomic distribution clearly resembled the distribution of mutations in the ageing mutation signature (C>T mutations in a lagging G context) and its complementary signature (T>C mutations in a lagging G context). This suggests that the loss of *CG* dinucleotides during evolution may be due to a similar mechanism to the ageing signature (spontaneous cytosine deamination in hypermethylated CpG islands). **e,** We compared the frequency of the substitution types that contained a *CG* motif in their trinucleotide context (yellow dots) against those that did not contain a *CG* motif (blue dots) for all classical trinucleotide mutation signatures. Scatter plots show that there was a strong suppression of *CG* dinucleotides for at least 10 trinucleotide signatures. Significance values were derived from Welch's t-test. **f-g,** We developed a Markov chain model to characterize the sequence composition of the human exome (Methods). Markov models assume that there is a constant *n* for which genomic positions with a distance larger than *n* do not interfere with each other's distributions (Markov order *n*). The major parameter of our Markov model is its maximal order, reflecting its context dependency. We tested the accuracy of Markov models with increasing orders (left to right) to predict the frequency of 6-nucleotide sequences in the reference genome. **f,** Q-Q-plots evaluate the calibration of our model for the sequence composition of the reference genome. The quantiles of the observed distribution (y-axis) are plotted against the quantiles of the expected distribution (x-axis) based on the Markov model. A close fit of the Q-Q plot to the diagonal (inflation factor $\lambda$ close to 1) suggests that the model is accurately calibrated. **g,** In parallel to the Q-Q-plots, we compared the observed and modeled frequency ratios (frequencies normalized against a uniform distribution) of nucleotide sequences in the human exome directly. Each dot represents an individual 6-nucleotide sequence. Expected frequency ratios (x-axis) are plotted against observed ratios (y-axis). Dot density is represented by dot colors, ranging from dark red (low density) to yellow (high density). In both analyses in (**f**) and (**g**), a Markov chain order of 2 nucleotides (16 Markov states) delivered the most precise model (orange). This finding suggests that underrepresentation of the *CG* motif was the main characteristic of the human exome, which had to be incorporated into the Markov model to capture the sequence composition of the human exome.

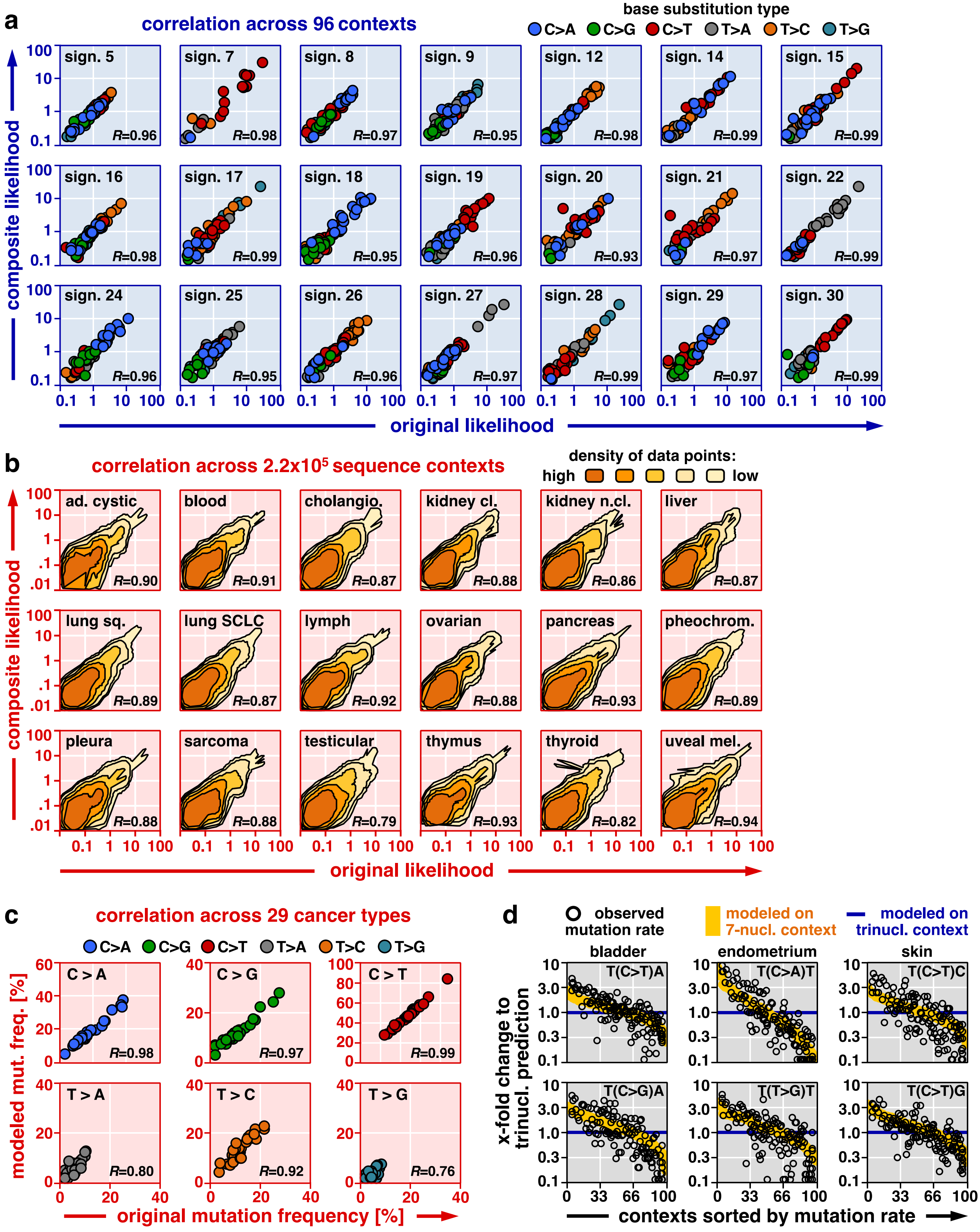


*(figure legend on next page)*

**Extended Data Fig. S7 | Modeling of context-specific mutation probabilities based on the broad nucleotide context.** This figure completes the plots shown in **Figure 1d** and **1f**. **a,** The original likelihood (x-axis) is plotted against the composite likelihood (y-axis) for each possible trinucleotide context. Dot colors indicate the base substitution types. Pearson correlations are annotated on the bottom right. **b,** Relative mutation counts (x-axis) are plotted against the expected mutation probabilities, derived from the composite likelihood (y-axis). As the number of possible 7-nucleotide sequence contexts was too large to be visualized directly, we visualized the data point density. The Pearson correlation of each plot is annotated on the bottom right. **c,** We plotted the original mutational likelihood (x-axis) against the composite likelihood (y-axis) for the 6 different base substitution types across the 29 cancer types examined in this study. Each dot represents an individual cancer type. The mutational likelihood of a mutation type was derived as the product of the mutational likelihood of its reference nucleotide (C vs. T) and its substitution type (type-I-transversions, type-II-transversions, transitions). For instance, for C>T substitutions the likelihood associated with C/G reference nucleotides was multiplied by the likelihood associated with transitions (C>T, G>A, T>C, A>G). The Pearson correlation of each plot is annotated on the bottom right.
**d,** We further examined whether flanking nucleotides outside of the immediate trinucleotide context had a substantial impact on the context-specific mutation probabilities in the composite likelihood model. For this purpose, we compared the mutation probabilities, modeled based on tri- (blue) and 7-nucleotide contexts (yellow), respectively, with the original likelihoods, derived from the mutation counts (black) shown in Figure 1f. Black circles indicate the ratio between the observed likelihoods and the corresponding trinucleotide-specific likelihoods (y-axis). Similarly, the orange line displays the ratio between the likelihoods, derived from the 7-nucleotide and trinucleotide contexts, respectively (y-axis). Data points are sorted according to the modeled mutation rates, derived from the 7-nucleotide context (x-axis). These plots demonstrate that local mutation probabilities are heterogeneous, even across positions with the same surrounding trinucleotide context (flanking 5' and 3' nucleotides). Accounting for flanking nucleotides outside of the trinucleotide context largely reduced this heterogeneity, indicating that these nucleotides had a substantial impact on the local context-specific mutation probability. Figure S8 provides a quantitative analysis of the signal outside of the trinucleotide context.

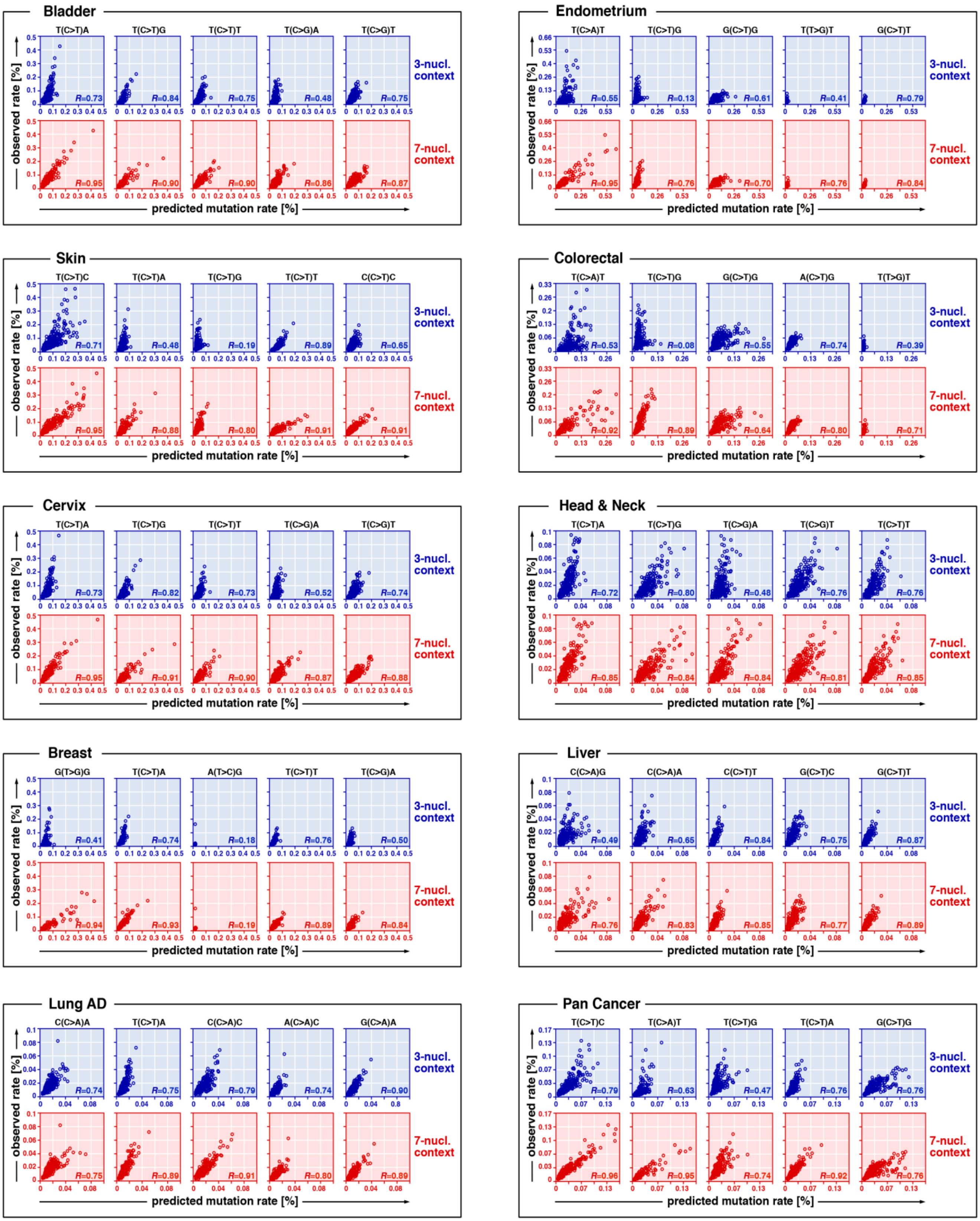


*(figure legend on next page)*

**Extended Data Fig. S8 | Quantification of the impact of flanking nucleotides outside of the trinucleotide context on local mutation probabilities in the composite likelihood.** Context-specific mutation probabilities are commonly modeled based on trinucleotide contexts (i.e., the flanking 5' and 3' nucleotides), whereas our composite likelihood model considered broad nucleotide contexts. We examined the impact of flanking nucleotides outside of the trinucleotide context on the mutation probability in the composite likelihood model. We counted for each possible 7-nucleotide context the number of mutations that fell into this specific 7-nucleotide context. We then normalized these counts against the total number of mutations, resulting in the observed mutation rate of each possible 7-nucleotide context (y-axis). We further derived the expected mutation rate of each possible 7-nucleotide from the composite likelihood model using either the full 7-nucleotide information (red, bottom, x-axis) or using the information of the inner trinucleotide context only (blue, top, x-axis). We plotted the predicted mutation rates (x-axis) against the observed mutation rates (y-axis) for 7-nucleotide contexts with the same inner trinucleotide context (annotated on the top). Each dot in these plots represents an individual 7-nucleotide context. The Pearson correlation of each plot is annotated on the bottom right. In contrast to the plots shown in Figure S7d, we compared the raw mutation counts, i.e. we did not normalize the raw mutation counts to their representation in the human reference exome.

We noticed that several of the trinucleotide correlation plots (blue) displayed more than one correlated groups of dots. For instance, plots in the upper rows of skin, endometrium and colorectal cancer display this phenomenon. Incorporating the nucleotides outside of the trinucleotide sequence context into the model accounts for this phenomenon (red plots), resulting a higher correlation between observed and predicted mutation rates. This observation reflects the impact of nucleotides outside of the trinucleotide sequence context on the raw mutation counts of individual genomic positions (blue plots). We thus concluded that incorporating the flanking nucleotides outside of the trinucleotide context into the composite likelihood model refined our approximation of local mutation probabilities.

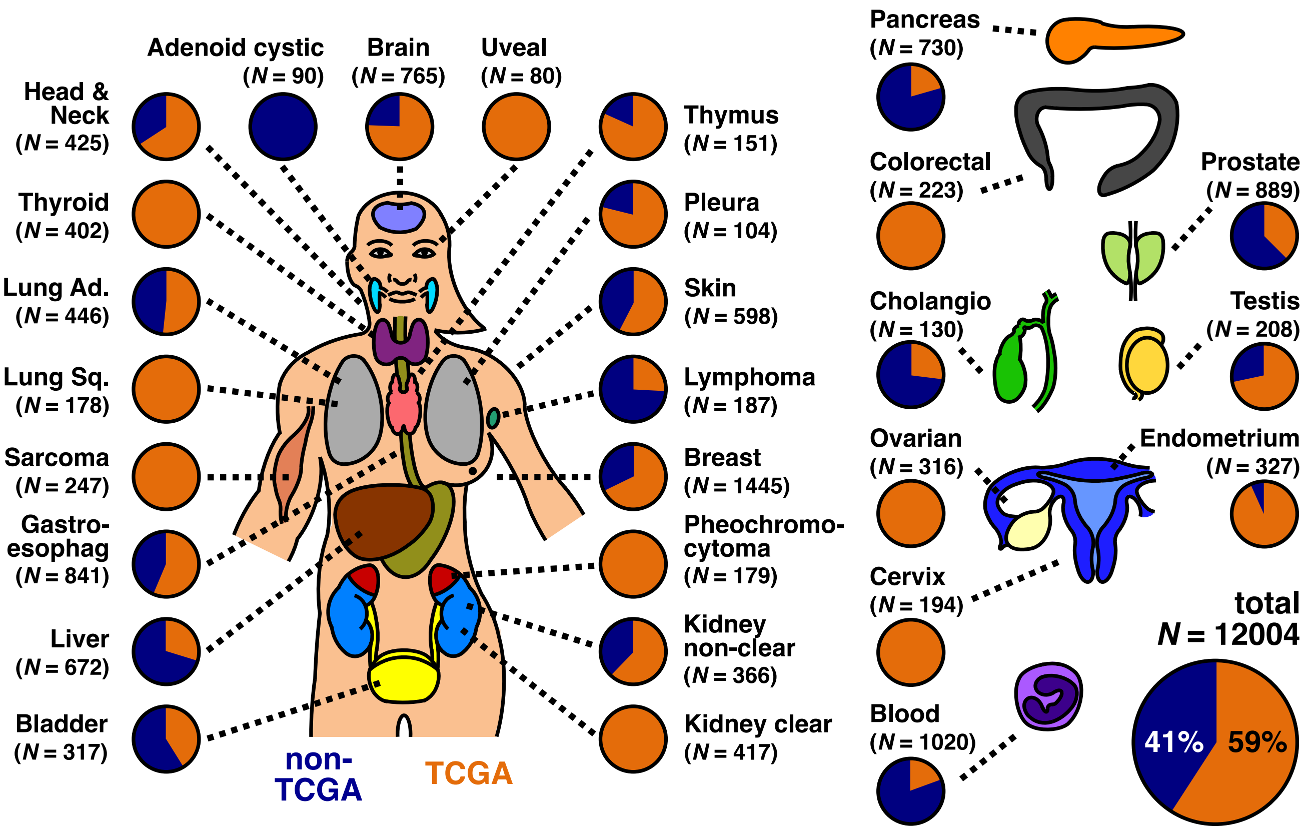


**Extended Data Fig. S9 | A large-scale cohort of whole-exome sequencing data to discover rare cancer genes.**
To discover novel candidate cancer genes, we analyzed sequencing data from 12,004 individual tumor samples using the statistical framework that we developed in this study (mutational recurrence and nucleotide context). Our study cohort contained whole-exome sequencing data from 32 TCGA-related (orange) and 57 TCGA-independent (blue) projects. A detailed characterization of the data sources included in this analysis can be found in Table S1.

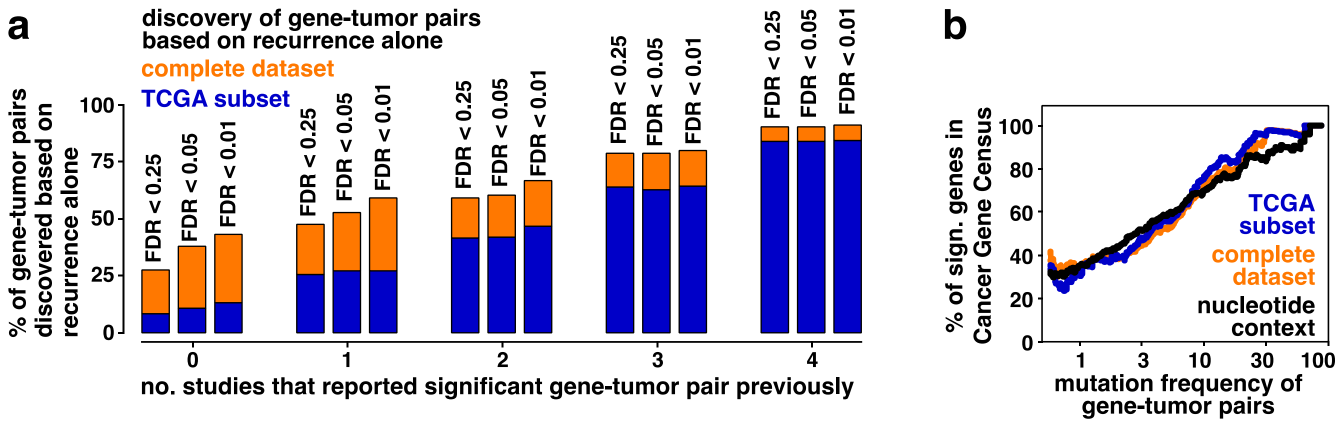


**Extended Data Fig. S10 | Discovery and characterization of new candidate cancer genes. a,** We counted for each significant gene-tumor pair gene in how many computational studies it had been reported as significantly mutated previously (x-axis). Further, we determined whether the gene-tumor pair was also discovered based on its recurrence alone using an established recurrence-based approach on our complete (orange) or TCGA (blue) dataset (y-axis). These plots show that the "re-discovery" rates shown in Figure 3c are stable for various false-discovery rate (FDR) thresholds. **b,** We counted how many significant gene-tumor pairs involved canonical cancer genes in the Cancer Gene Census, as a surrogate marker for true-positive findings. Gene-tumor pairs were either identified by recurrence (blue: TCGA dataset, orange: full dataset) or by considering recurrence and nucleotide context (black). We plotted the fraction of gene-tumor pairs in the Cancer Gene Census (y-axis) against their mutation frequency (x-axis). Considering the nucleotide context yielded a higher number of gene-tumor pairs with mutation frequencies <5% (Fig. 3d). However, after correcting for mutation frequencies, our statistical framework and the established recurrence-based approach discovered similar fractions of canonical cancer genes, suggesting that the false-discovery rate of our approach does not exceed the false-discovery rates of established recurrence-based approaches.

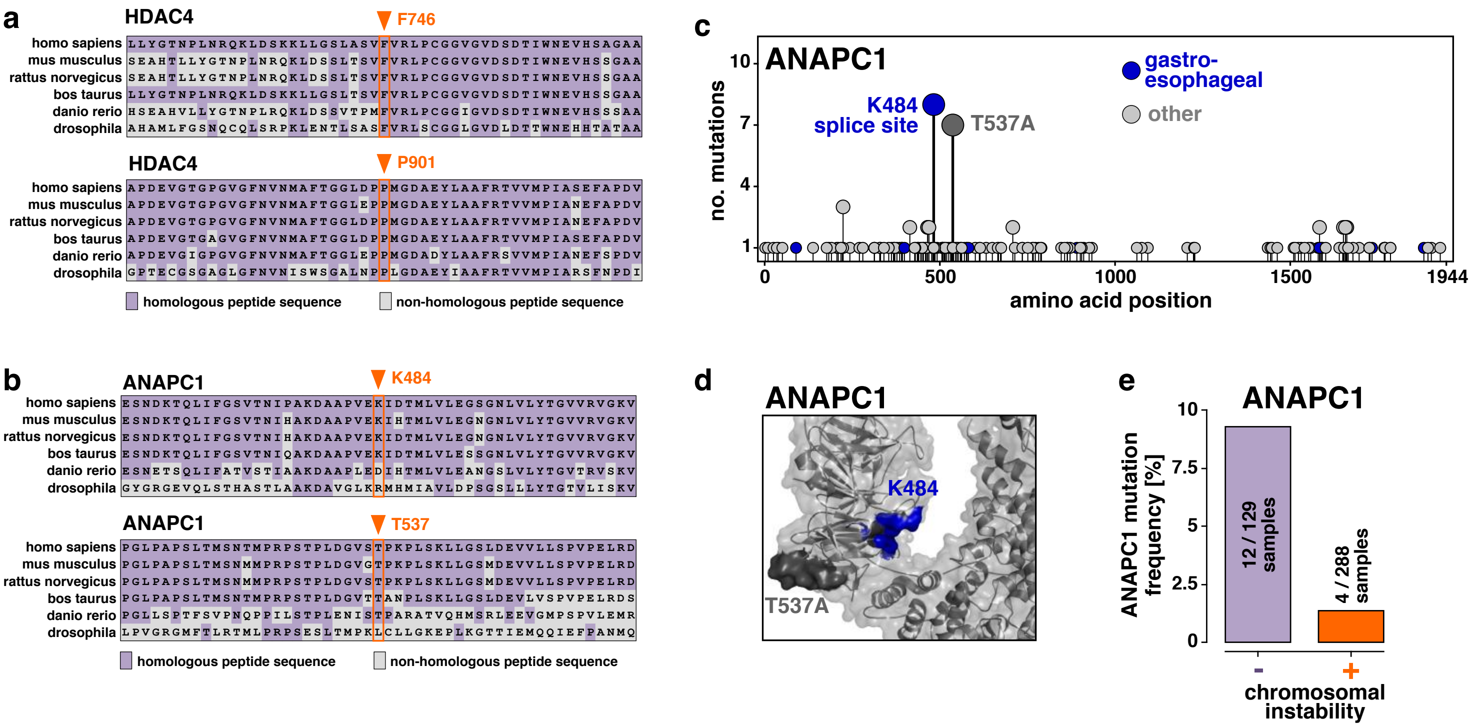


**Extended Data Fig. S11 | Characterization of the candidate cancer gene *ANAPC1*. a-b,** We aligned the peptide sequences of *HDAC4* (**a**) and *ANAPC1* (**b**) across six species (rows) around the positions (orange) that were recurrently altered by somatic mutations. Homologous amino acids are highlighted in violet; non-homologous amino acids are colored in gray. These alignments suggest that mutational hotspots in *HDAC4* and *ANAPC1* target evolutionarily conserved protein domains. **c-e,** Exemplary evidence for the candidate cancer gene *ANAPC1*, which was discovered as a significantly mutated gene in gastroesophageal cancer by considering nucleotide context and recurrence (FDR = 0.029) but not based on recurrence alone (FDR=1.0). (**c**) The distribution of *ANAPC1* mutations is visualized as a needle plot. For each amino acid substitution the number of samples (y-axis) is plotted against its amino acid position in the peptide sequence (x-axis). Dot colors reflect the tumor types (blue: gastroesophageal cancer, gray: other tumor types) in which the amino acid substitution was detected. (**d**) The position of the two mutational hotspots was visualized using a previously published crystal structure (PDB: 5G05). It has been reported previously that Cdk1-mediated phosphorylation of threonine 537 (T537) in ANAPC1 increases the catalytic activity of this complex by ~6-fold. Substitution of this T537 residue, which is located in the 500s loop of the WD40 repeat domain, by the nonpolar amino acid alanine may decrease the activity of the anaphase-promoting complex and thus prolong the transition from the metaphase to the anaphase. **e,** A recent study suggested that a prolonging the transition from the metaphase to the anaphase limits excessive chromosomal instability (CIN) in cancer. Using the CIN annotations from the TCGA gastroesophagal marker paper we investigated this hypothesis. In concordance with the functional results reported previously, we found that mutations in *ANAPC1* were negatively associated with chromosomal instability in our study cohort.

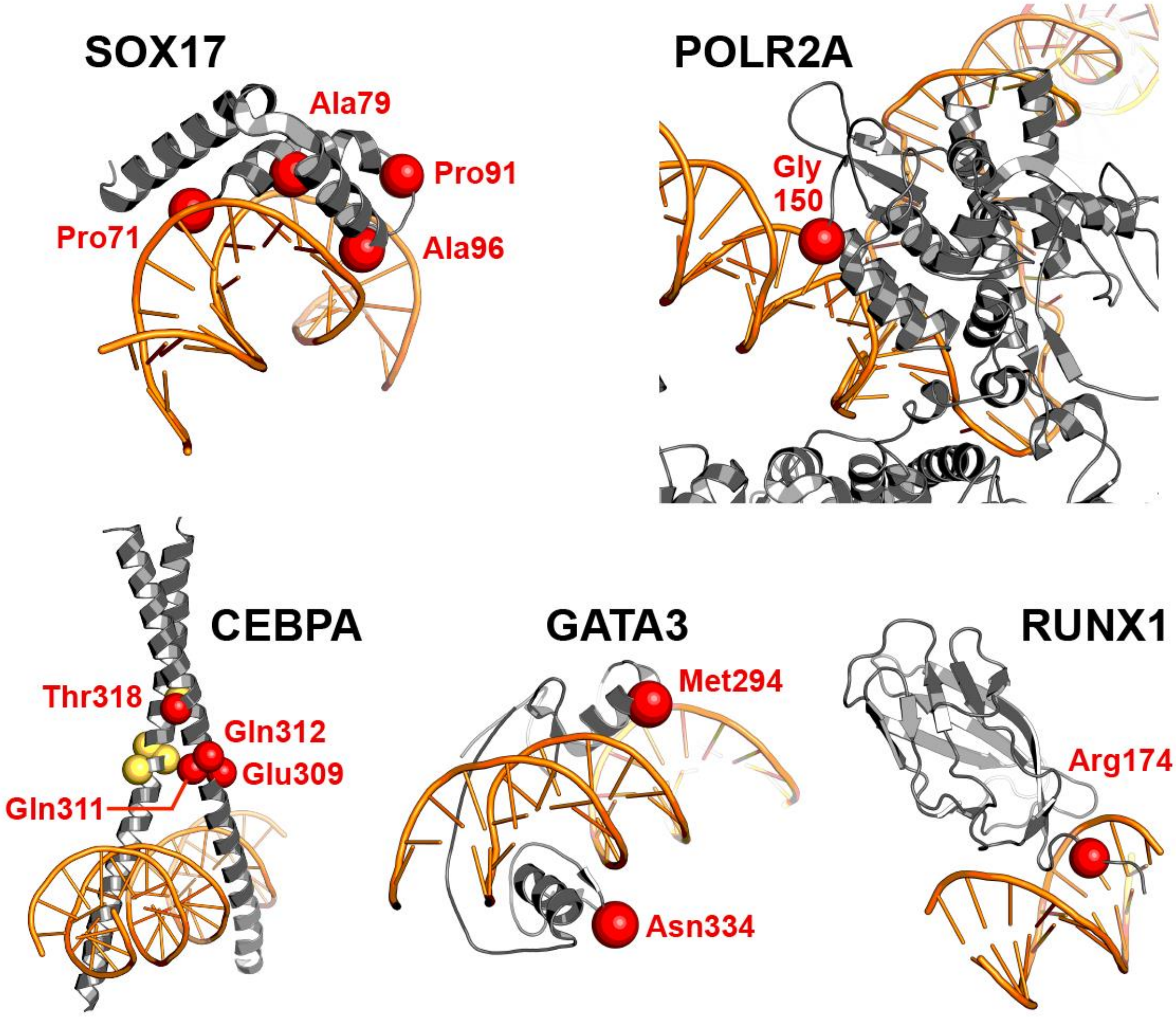


**Extended Data Fig. S12 | Visualization of recurrent mutations in positions involved in the protein-DNA interaction.** Using previously published crystal structures we noticed that the transcription factors *SOX17* (PDB: 4A3N superposed with 3F27), *CEBPA* (PDB: 1NWQ), *GATA3* (PDB: 4HCA), and *RUNX1* (PDB: 1H9D), as well as the RNA polymerase *POLR2A* (PDB: 5IYB) harbored recurrent mutations in positions that were relevant for the protein-DNA interaction. More precisely, we identified ***SOX17*** as a significantly mutated gene (FDR = 0.01) in endometrial cancer. *SOX17* mutations are recurrently located in the high-mobility-group box domain (HMG domain) of SOX17 at the SOX17-DNA interface. This structural motif is involved in binding and bending DNA in the minor grove in order to induce the assembly transcriptional machineries. ***POLR2A*** was identified as a significantly mutated gene in lung adenocarcinomas (FDR = $1.07\times10^{-5}$). The open complex of a cryo-EM multicomponent structure where the melted single-stranded template DNA is inserted into the active site and RNA polymerase II locates the transcription start site is visualized. *POLR2A* harbors recurrent mutations at the end of an alpha helical segment that in this state is directly pointed at the major groove of the double stranded DNA alongside the active site of replication. ***CEBPA*** was identified as significantly mutated gene (FDR = $4.58\times10^{-7}$) in hematological malignancies. *CEBPA* harbors recurrent mutations at the cross-over interface of the two CEBPA homodimers constituting the active transcription factor homodimer. ***GATA3*** was identified as a significantly mutated gene (FDR < $10^{-20}$) in breast cancer. The residue Asn334 is located in the GATA-type 2 zinc finger (res317-res341), whereas the residue Met294 is located peripheral to the GATA-type 1 zinc finger (res263-res287) domain pointing towards the major groove of the DNA binding partner. Finally, ***RUNX1*** was identified as a significantly mutated gene in breast cancer (FDR = $1.99\times10^{-4}$) and hematological malignancies (FDR = $7.96\times10^{-5}$). The residue Arg174 plays an important role for the DNA recognition mechanism as it facilitates the formation of hydrogen bond interactions to a guanosine base from the consensus DNA binding sequence of RUNX1. Mutational studies demonstrated that the R174A mutation (and others adjacent to Arg174) leads to severely perturbed DNA binding.

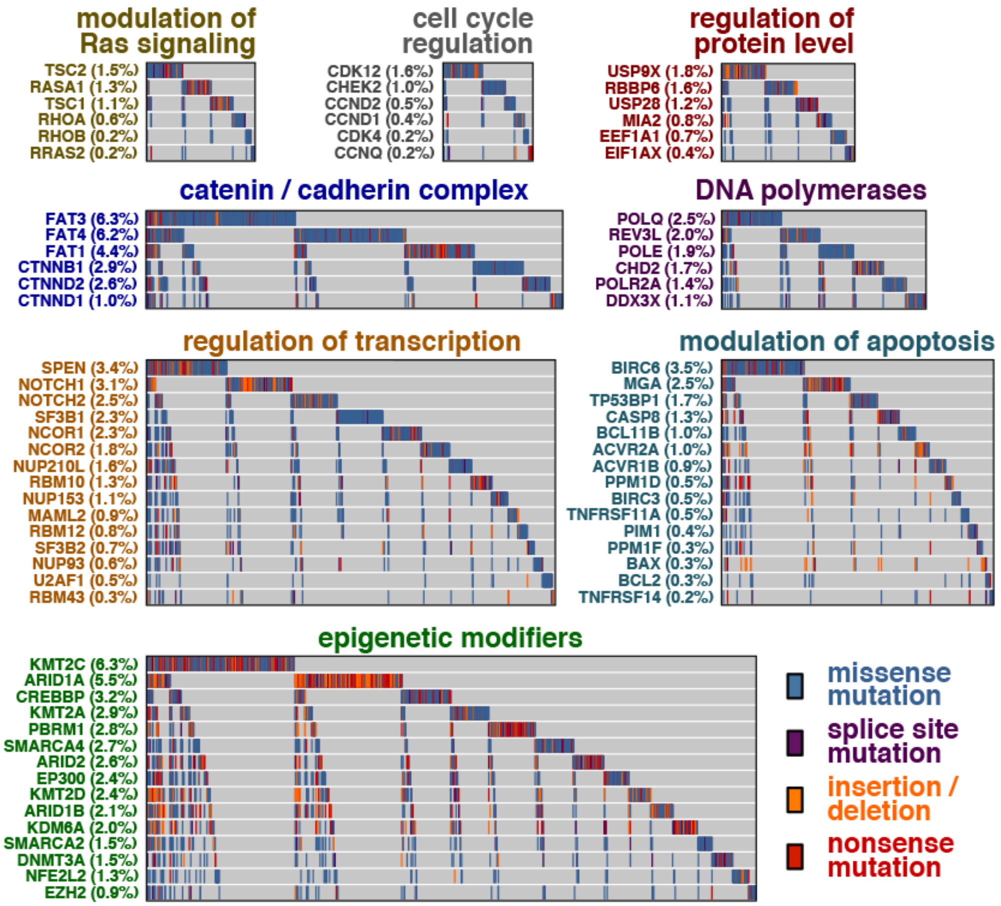


**Extended Data Fig. S13 | Novel gene-tumor pairs in cancer-relevant signaling complexes.** Several novel gene-tumor pairs (i.e., new cancer genes or known cancer genes in a new tumor type context) fell into one of the eight signaling complexes displayed in this figure. For each signaling complex, the distribution of somatic mutations is visualized (x-axis: mutant samples, y-axis: significantly mutant genes, colors: mutation category). Due to their low overall mutation frequencies, the relevance of these pathways might have remained undervalued in previous computational studies using recurrence-based approaches for cancer gene discovery. In aggregate, however, mutations in these signaling pathways may be functionally relevant for a substantial number of tumor patients.

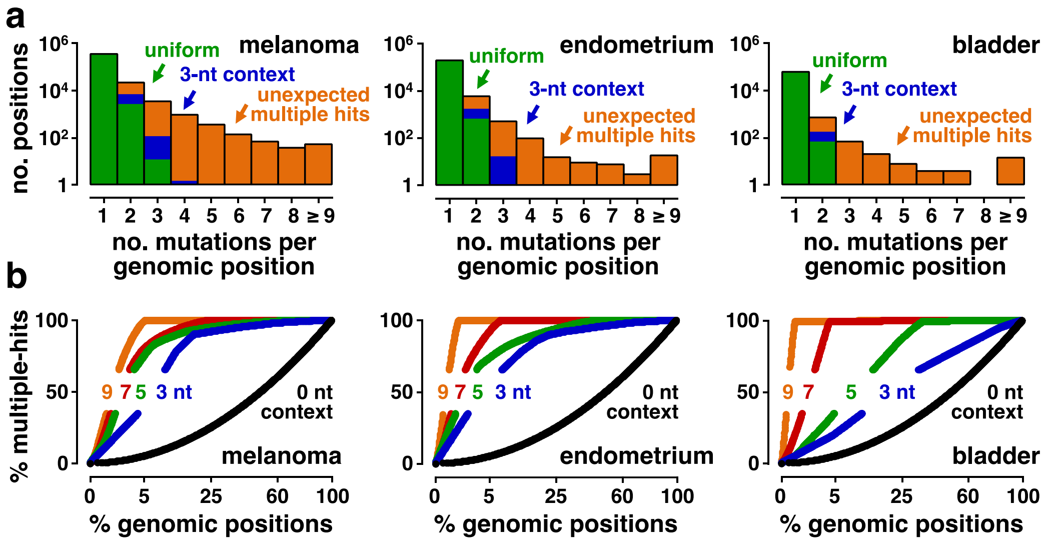


**Extended Data Fig. S14 | Passenger mutations display a conglomerative distribution pattern in the exomes of human cancer.** Data relate to the genomic distribution of mutations in 598 melanomas (left), 327 endometrial carcinomas (middle) and 317 bladder cancers (right). **a,** We counted the number of genomic positions that were hit by more than one independent mutation. The histograms display the number of genomic positions in the exome, containing 1, 2, … , 8 or ≥9 independent mutations, respectively (orange: observed histogram; green: histogram expected based on a uniform distribution; blue: histogram, obtained by redistributing mutations conditional on their trinucleotide context). To distinguish between random accumulations of passenger mutations and true mutational hotspots, accurate modeling of this conglomerative distribution pattern is required.

**b,** We asked whether integration of flanking nucleotides outside of the trinucleotide context might help understand the conglomerative distribution pattern of passenger mutations. We determined the probability of each genomic position in the exome of being hit by more than one mutation, based on the number of nucleotides (nt) incorporated into the surrounding sequence context. We then sorted the genomic positions in descending order of those probabilities (x-axis, logarithmic). The genomic positions are plotted against the cumulative fraction of multiple hit positions (y-axis, positions which contain more than one mutation). A uniform distribution of mutations served as a negative control (black). These data suggest that considering the broad nucleotide context is particularly important to identify genomic positions with exceptionally high mutation probabilities, which is required to inform mutational hotspot discovery.

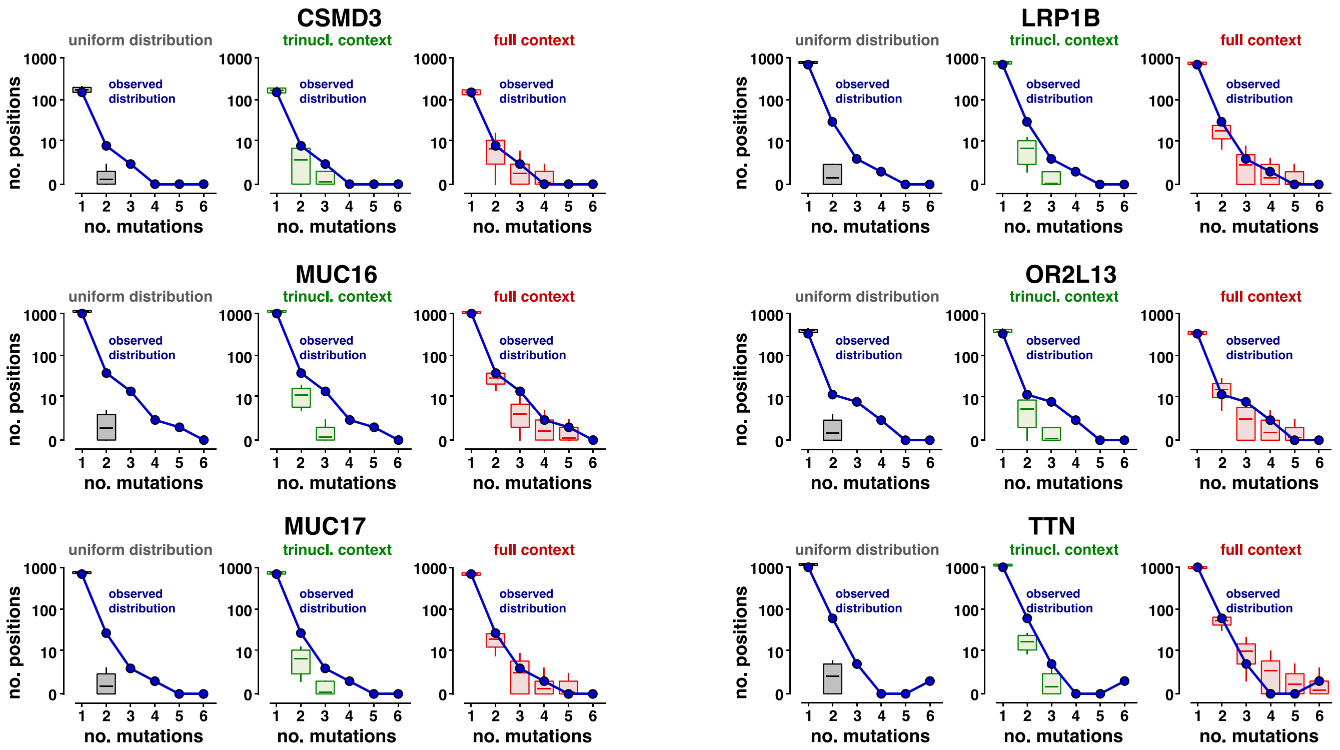


**Extended Data Fig. S15 | Modeling the distribution of passenger mutations based on nucleotide context.** A major prerequisite for the discovery of novel cancer genes was our model sufficiently calibrated to the background distribution. Hence, we examined whether our model was accurately calibrated to the distribution of passenger mutations in non-cancer-related genes that had been reported as false-positive findings in previous studies (*CSMD3*, *MUC16*, *MUC17*, *LRP1B*, *OR2L13*, *TTN* in melanoma). We counted the number of positions that were hit by more than one mutation (blue). We then randomly re-distributed mutations in these genes (10,000 iterations), assuming a uniform distribution (gray), trinucleotide-specific mutation probabilities (green), or mutation probabilities derived from the full composite likelihood model (red). In each iteration, we counted the number of positions containing 1, … , 5, and ≥6 mutations, respectively. The distributions of these simulated counts are represented as box plots. Boxes mark the interquartile range; vertical lines denote the 5%-95% percentile range; horizontal lines mark the group medians. These experiments revealed that the uniform model (gray) and the trinucleotide context model (green) systematically underestimated the number of positions that were hit by more than one mutation. Considering the broad nucleotide context (red) provided a closer approximation ofthe local distribution of passenger mutations in these hypermutant non-cancer-associated genes.

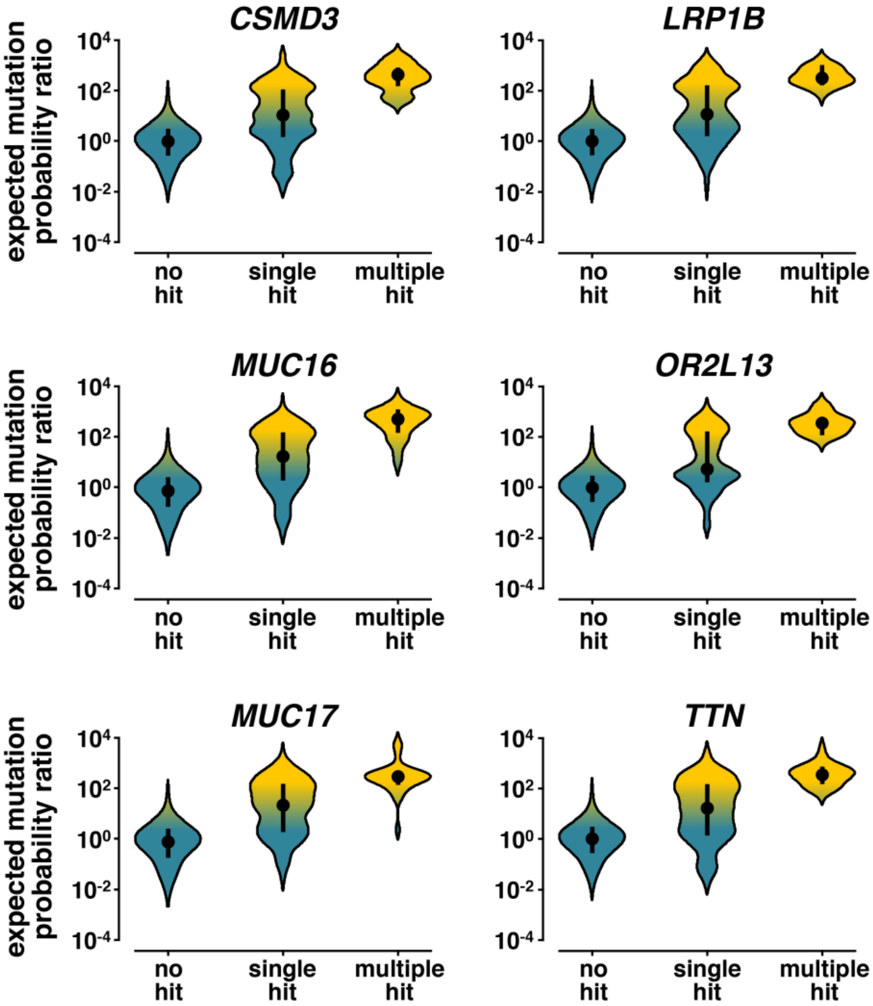


**Extended Data Fig. S16 | Accounting for the broad nucleotide context identifies genomic positions in human cancer that are prone to accumulate multiple passenger mutations.** We asked whether considering the broad nucleotide context informed the discovery of mutational hotspots in cancer. Critically, random accumulations of passenger mutations in positions with high mutation probabilities need to be distinguished from mutational hotspots. For this purpose, we examined the distribution of the mutation probability ratios derived from our composite likelihood model in six non-cancer-related genes with high background mutation rates (*CSMD3*, *LRP1B*, *MUC16*, *OR2L13*, *MUC17*, *TTN* in melanoma). Violin plots display the distribution of the mutational likelihood in individual positions, depending on whether they contained no mutations (no hit), a single mutation (single hit) or which contained more than one mutation (multiple hit). A mutational likelihood >1 indicates that the position is expected to contain more mutations than based on a uniform distribution. Probability ratios of the non-mutant positions stably varied around 1, and there was ~100× increase in the mutation probability ratios for multiple-hit positions. Positions with a single mutation displayed a bimodal distribution inbetween. This finding suggests that our context-dependent composite likelihood model accurately identifies positions, which have a high probability of accumulating somatic passenger mutations. Thus, considering the broad nucleotide context informs discovery of mutational cancer hotspots and cancer driver mutations.

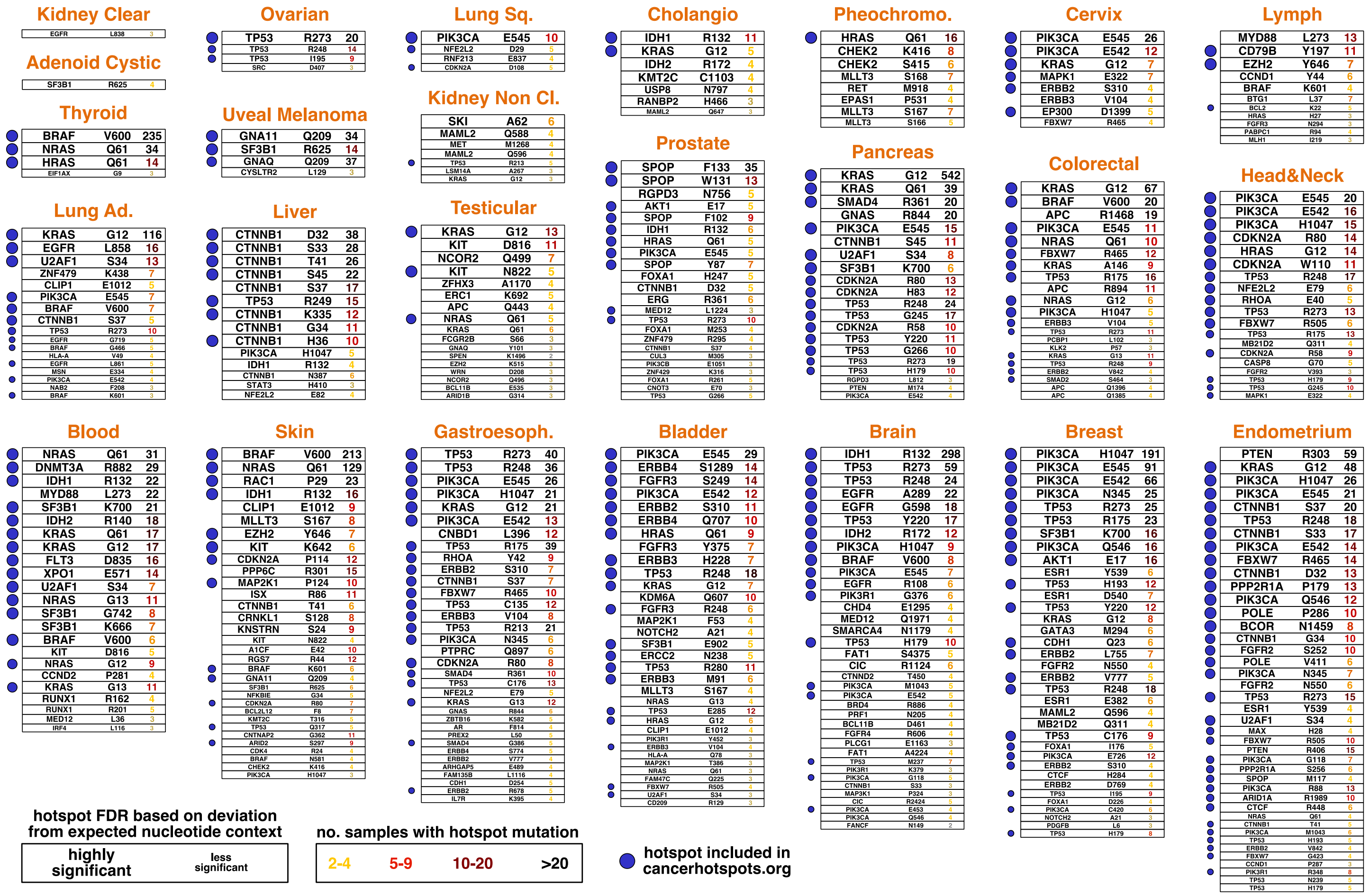


**Extended Data Fig. S17 | Discovery of mutational hotspots in cancer.** We examined the feasibility of the nucleotide context to distinguish between spontaneous accumulations of passenger mutations. For this purpose, we counted for each genomic position its number of mutations and determined whether this mutation count exceeded our expectation based on its context-dependent mutation probability. Based on this comparison between the mutation count and the corresponding mutation probability, we derived a significance value for each position that contained at least 2 mutations. We corrected these significance values for multiple hypothesis testing (false-discovery rates). Positions that exceeded the mutation probability significantly are listed in descending order according to their number of mutations. Further, we annotated for each significant position the gene name as well as the amino acid residue. Font sizes indicate hotspot significance values; font colors reflect mutation counts. Blue dots indicate whether the hotspot had been reported in two large-scale sophisticated hotspot analyses (data from cancerhotspots.org). These analyses suggest that characterization of the broad nucleotide context informs the discovery of mutational hotspots. Details on the statistics that we used for hotspot discovery can be found in the Methods. We note that this statistics is part of the full statistical framework that we used for cancer gene discovery and contributes to the significance values shown in Figure 4.

**Extended Data Table S1 | Sources of sequencing data included in this study.** This table contains literature references to all 89 studies, from which sequencing data were included in this study; 32 of these sequencing projects are TCGA-related. This table further annotates how many samples from each study are contained in our study cohort.

**Extended Data Table S2 | Stratification of gene-tumor pairs based on their literature support.** We stratified the significant gene-tumor pairs (i.e., pairs of significantly mutated genes and their associated cancer type) by their literature support. As a first layer, we examined whether the gene-tumor pair had been consistently reported in at least two previous computational studies (blue: known cancer genes). As a second layer, we searched for publications with experimental or clinical data that functionally implicated our findings in the same tumor types in which we discovered them as significantly mutated (red: literature level A). As a third layer, we searched for experimental data that functionally implicated the gene in cancer, but not in the same tumor type (orange: literature level B). Genes without any literature support are colored in gray. The table further lists for each gene-tumor pair its gene name (official symbol of the NCBI RefSeq database), its cancer type, its full gene name, its validation level, whether the gene was part of the Cancer Gene Census, its mutation frequency, its significance value (FDR based on recurrence and nucleotide context), and its recurrence-based significance (FDR derived from an established recurrence-based approach).

**Extended Data Table S3 | Stratification of genes based on their literature support.** In parallel to Table S2, we stratified significantly mutated genes by their literature support. This table further lists for each gene its gene name (official symbol of the NCBI RefSeq database), its full name, the number of cancer types in which we detected it as significantly mutated, its validation level, whether the gene was part of the Cancer Gene Census, its maximal mutation frequency, its best significance value across all cancer types (FDR based on recurrence and nucleotide context), and its best recurrence-based significance (FDR derived from an established recurrence-based approach).
