## Supplementary material for "Discovery of cancer driver genes based on nucleotide context"

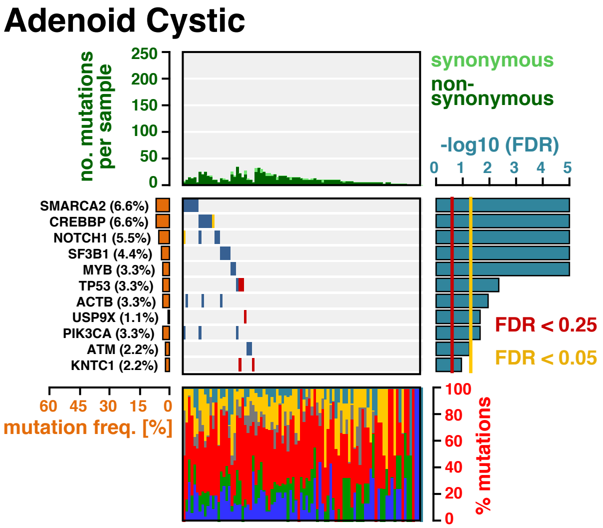


**Extended Data Fig. S18 | The landscape of driver mutations in adenoid cystic carcinomas.** The color-coded matrix in the center displays the distribution of non-synonymous mutations across individual tumor samples (x-axis) in the significantly mutant genes, identified by MutPanning (y-axis). The bar graph on top shows the total number of mutations in each sample. Non-synonymous mutations are colored in dark green; synonymous mutations are colored in light green. The relative contribution of the six mutation types (blue: C>A, green: C>G, red: C>T, gray: T>A, yellow: T>C, cyan: T>G) to the mutational burden of each sample is shown in the bar graph below. The cancer type specific mutation frequency of each significant gene is shown on the left (orange). The False-discovery rates, computed by MutPanning (FDR, q-values), are displayed on the right. The red line indicates a cutoff at q<0.25; the yellow line indicates a threshold at q<0.05.

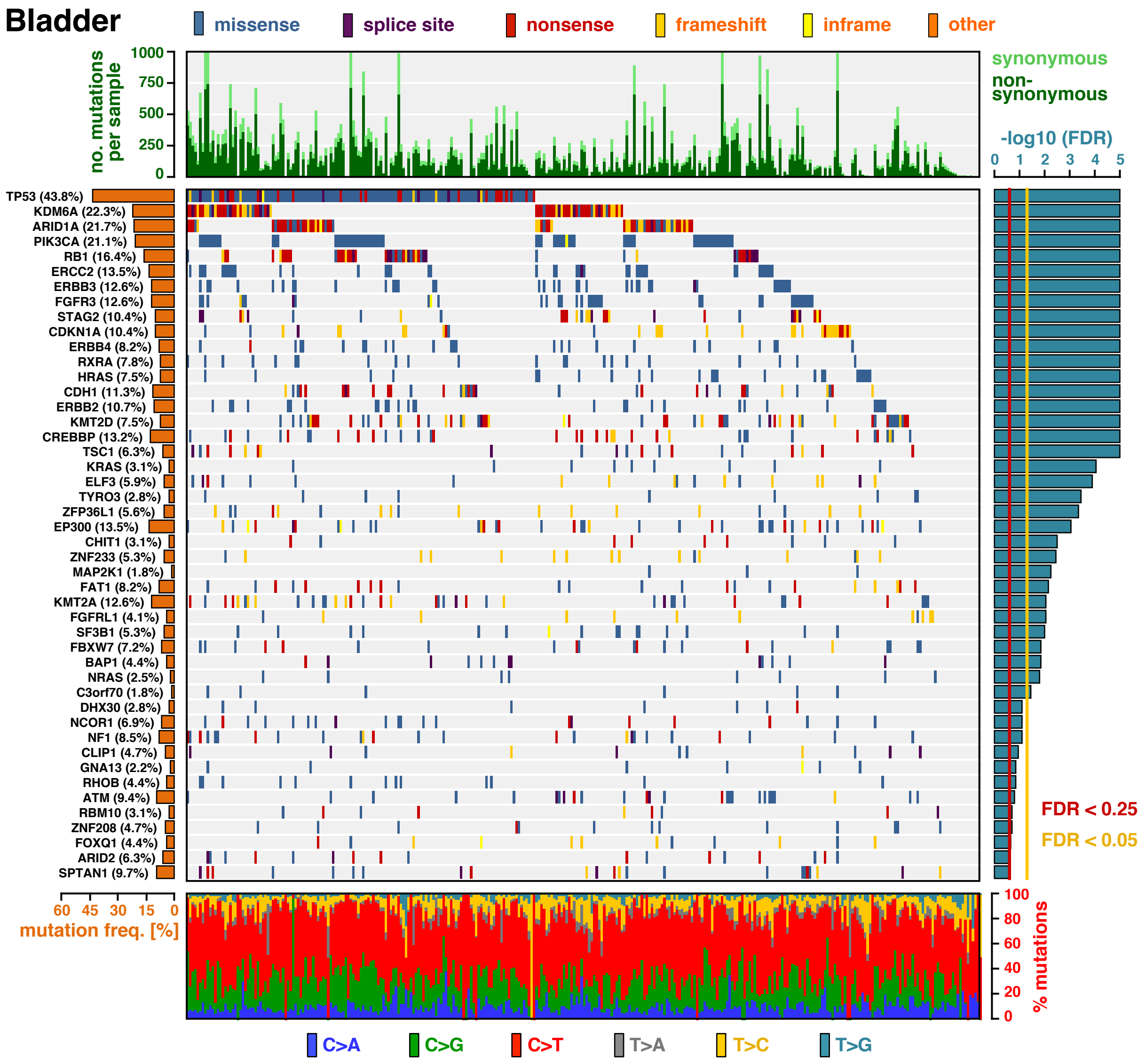


**Extended Data Fig. S19 | The landscape of driver mutations in bladder cancer.** The color-coded matrix in the center displays the distribution of non-synonymous mutations across individual tumor samples (x-axis) in the significantly mutant genes, identified by MutPanning (y-axis). The bar graph on top shows the total number of mutations in each sample. Non-synonymous mutations are colored in dark green; synonymous mutations are colored in light green. The relative contribution of the six mutation types (blue: C>A, green: C>G, red: C>T, gray: T>A, yellow: T>C, cyan: T>G) to the mutational burden of each sample is shown in the bar graph below. The cancer type specific mutation frequency of each significant gene is shown on the left (orange). The False-discovery rates, computed by MutPanning (FDR, q-values), are displayed on the right. The red line indicates a cutoff at q<0.25; the yellow line indicates a threshold at q<0.05.

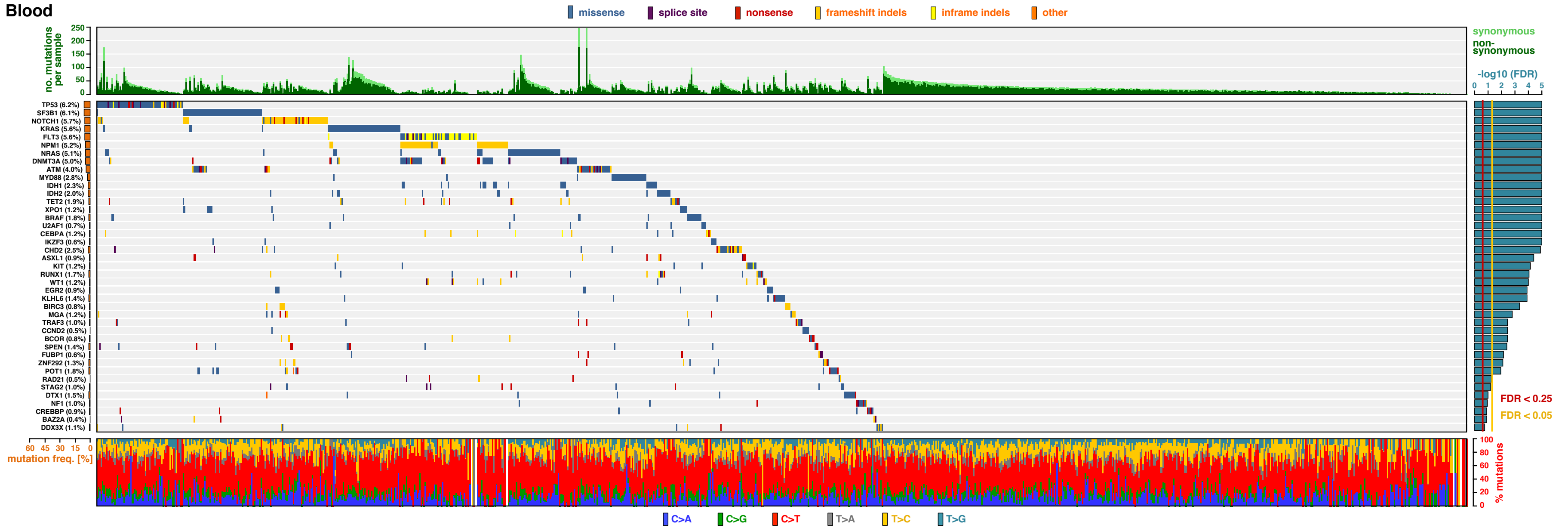


**Extended Data Fig. S20 | The landscape of driver mutations in hematological malignancies.** The color-coded matrix in the center displays the distribution of non-synonymous mutations across individual tumor samples (x-axis) in the significantly mutant genes, identified by MutPanning (y-axis). The bar graph on top shows the total number of mutations in each sample. Non-synonymous mutations are colored in dark green; synonymous mutations are colored in light green. The relative contribution of the six mutation types (blue: C>A, green: C>G, red: C>T, gray: T>A, yellow: T>C, cyan: T>G) to the mutational burden of each sample is shown in the bar graph below. The cancer type specific mutation frequency of each significant gene is shown on the left (orange). The False-discovery rates, computed by MutPanning (FDR, q-values), are displayed on the right. The red line indicates a cutoff at q<0.25; the yellow line indicates a threshold at q<0.05.

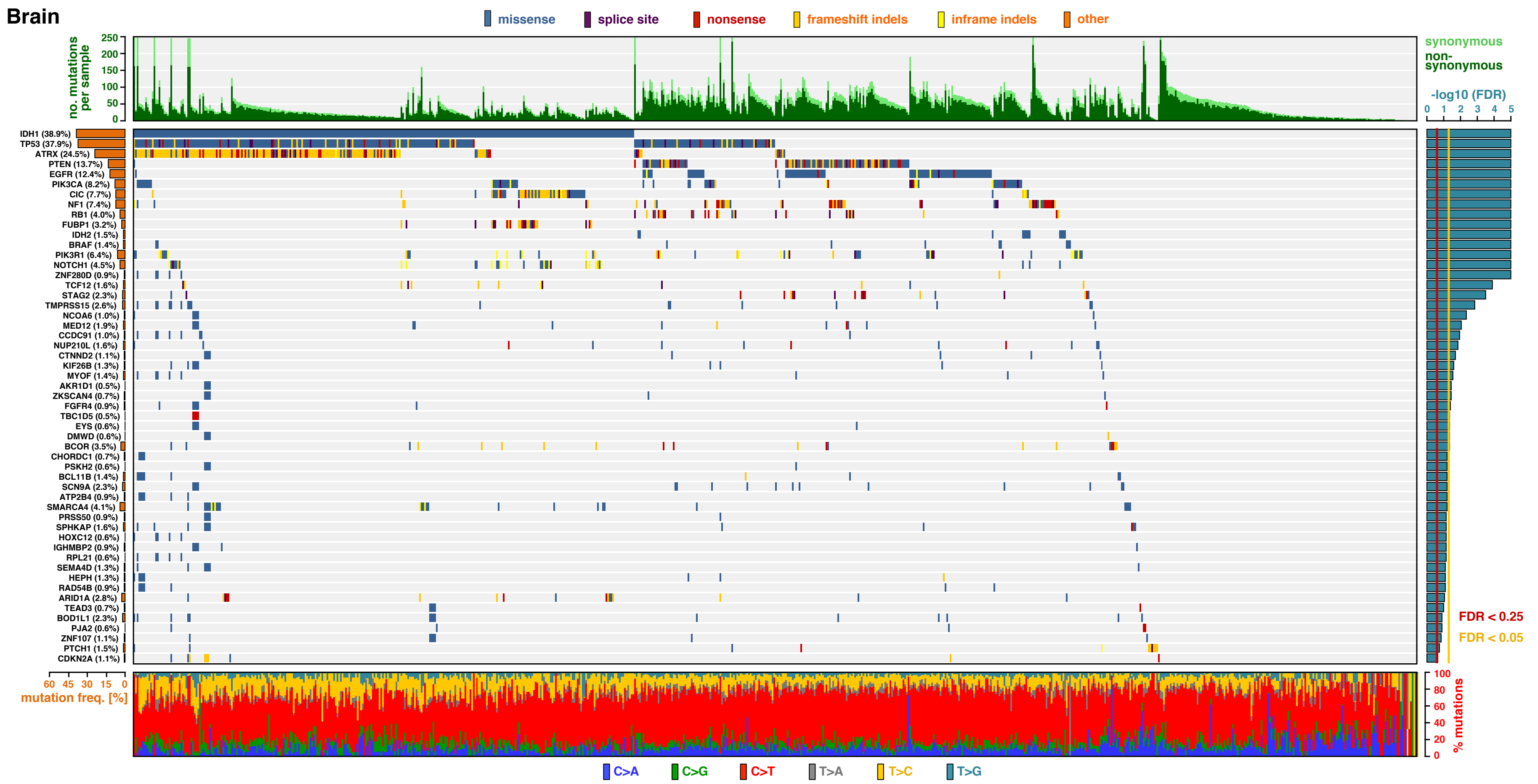


**Extended Data Fig. S21 | The landscape of driver mutations in brain tumors.** The color-coded matrix in the center displays the distribution of non-synonymous mutations across individual tumor samples (x-axis) in the significantly mutant genes, identified by MutPanning (y-axis). The bar graph on top shows the total number of mutations in each sample. Non-synonymous mutations are colored in dark green; synonymous mutations are colored in light green. The relative contribution of the six mutation types (blue: C>A, green: C>G, red: C>T, gray: T>A, yellow: T>C, cyan: T>G) to the mutational burden of each sample is shown in the bar graph below. The cancer type specific mutation frequency of each significant gene is shown on the left (orange). The False-discovery rates, computed by MutPanning (FDR, q-values), are displayed on the right. The red line indicates a cutoff at q<0.25; the yellow line indicates a threshold at q<0.05.

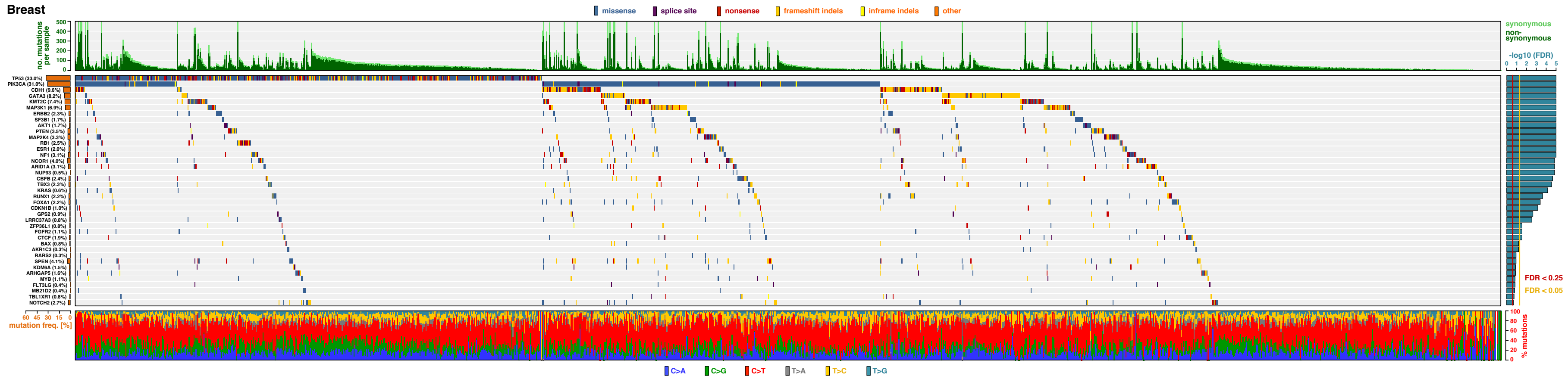


**Extended Data Fig. S22 | The landscape of driver mutations in breast cancer.** The color-coded matrix in the center displays the distribution of non-synonymous mutations across individual tumor samples (x-axis) in the significantly mutant genes, identified by MutPanning (y-axis). The bar graph on top shows the total number of mutations in each sample. Non-synonymous mutations are colored in dark green; synonymous mutations are colored in light green. The relative contribution of the six mutation types (blue: C>A, green: C>G, red: C>T, gray: T>A, yellow: T>C, cyan: T>G) to the mutational burden of each sample is shown in the bar graph below. The cancer type specific mutation frequency of each significant gene is shown on the left (orange). The False-discovery rates, computed by MutPanning (FDR, q-values), are displayed on the right. The red line indicates a cutoff at q<0.25; the yellow line indicates a threshold at q<0.05.

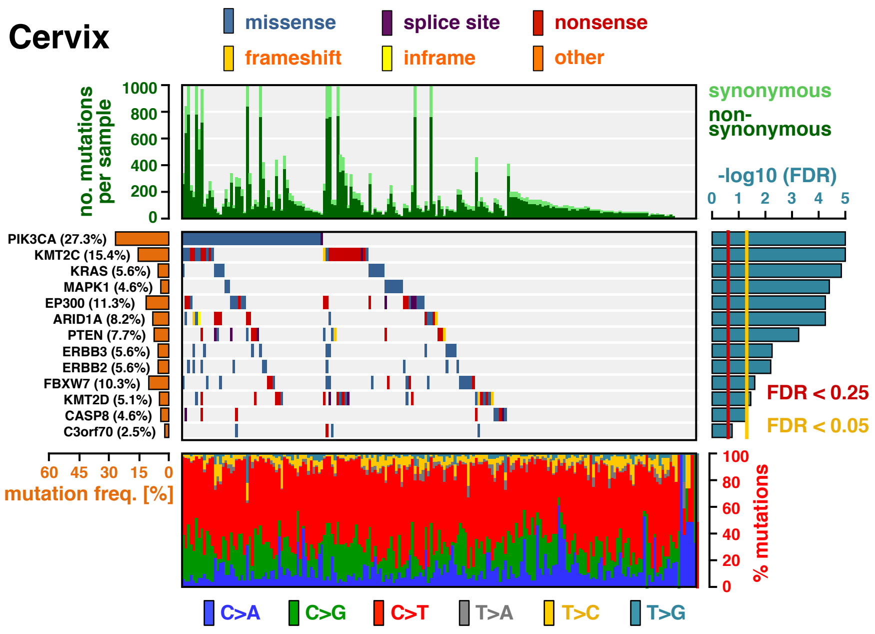


**Extended Data Fig. S23 | The landscape of driver mutations in cervical cancer.** The color-coded matrix in the center displays the distribution of non-synonymous mutations across individual tumor samples (x-axis) in the significantly mutant genes, identified by MutPanning (y-axis). The bar graph on top shows the total number of mutations in each sample. Non-synonymous mutations are colored in dark green; synonymous mutations are colored in light green. The relative contribution of the six mutation types (blue: C>A, green: C>G, red: C>T, gray: T>A, yellow: T>C, cyan: T>G) to the mutational burden of each sample is shown in the bar graph below. The cancer type specific mutation frequency of each significant gene is shown on the left (orange). The False-discovery rates, computed by MutPanning (FDR, q-values), are displayed on the right. The red line indicates a cutoff at q<0.25; the yellow line indicates a threshold at q<0.05.

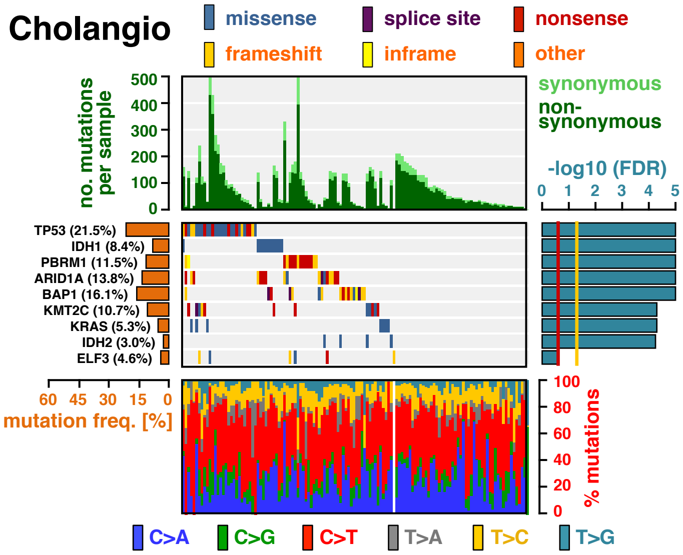


**Extended Data Fig. S24 | The landscape of driver mutations in cholangiocarcinoma.** The color-coded matrix in the center displays the distribution of non-synonymous mutations across individual tumor samples (x-axis) in the significantly mutant genes, identified by MutPanning (y-axis). The bar graph on top shows the total number of mutations in each sample. Non-synonymous mutations are colored in dark green; synonymous mutations are colored in light green. The relative contribution of the six mutation types (blue: C>A, green: C>G, red: C>T, gray: T>A, yellow: T>C, cyan: T>G) to the mutational burden of each sample is shown in the bar graph below. The cancer type specific mutation frequency of each significant gene is shown on the left (orange). The False-discovery rates, computed by MutPanning (FDR, q-values), are displayed on the right. The red line indicates a cutoff at q<0.25; the yellow line indicates a threshold at q<0.05.

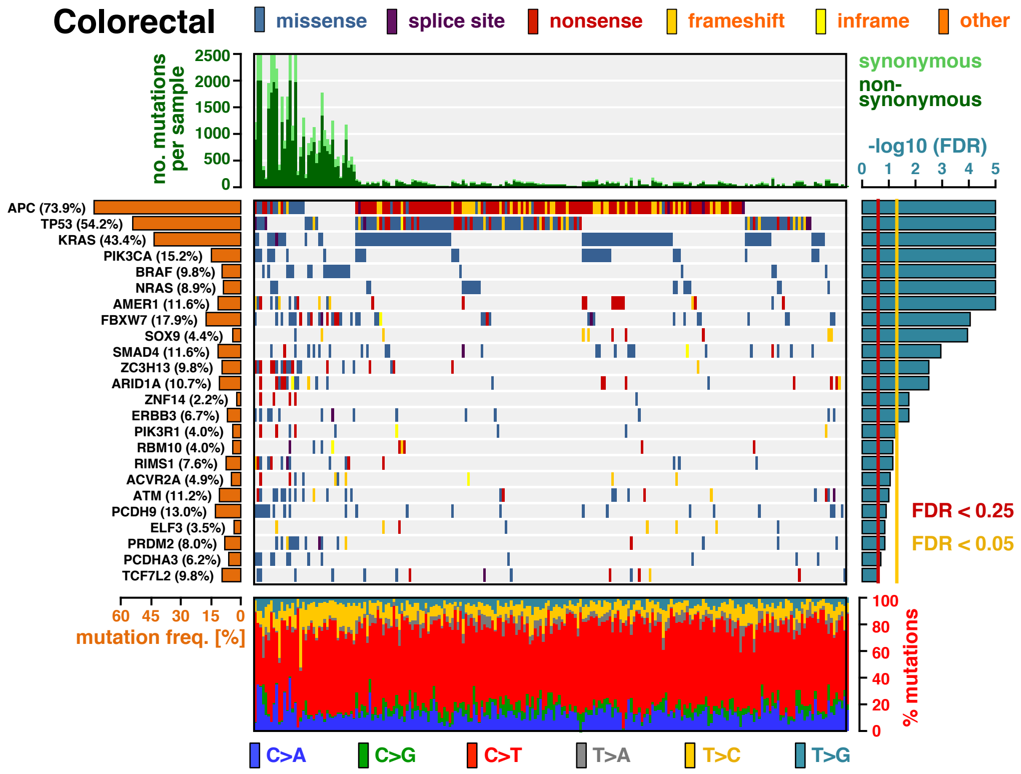


**Extended Data Fig. S25 | The landscape of driver mutations in colorectal cancer.** The color-coded matrix in the center displays the distribution of non-synonymous mutations across individual tumor samples (x-axis) in the significantly mutant genes, identified by MutPanning (y-axis). The bar graph on top shows the total number of mutations in each sample. Non-synonymous mutations are colored in dark green; synonymous mutations are colored in light green. The relative contribution of the six mutation types (blue: C>A, green: C>G, red: C>T, gray: T>A, yellow: T>C, cyan: T>G) to the mutational burden of each sample is shown in the bar graph below. The cancer type specific mutation frequency of each significant gene is shown on the left (orange). The False-discovery rates, computed by MutPanning (FDR, q-values), are displayed on the right. The red line indicates a cutoff at q<0.25; the yellow line indicates a threshold at q<0.05.

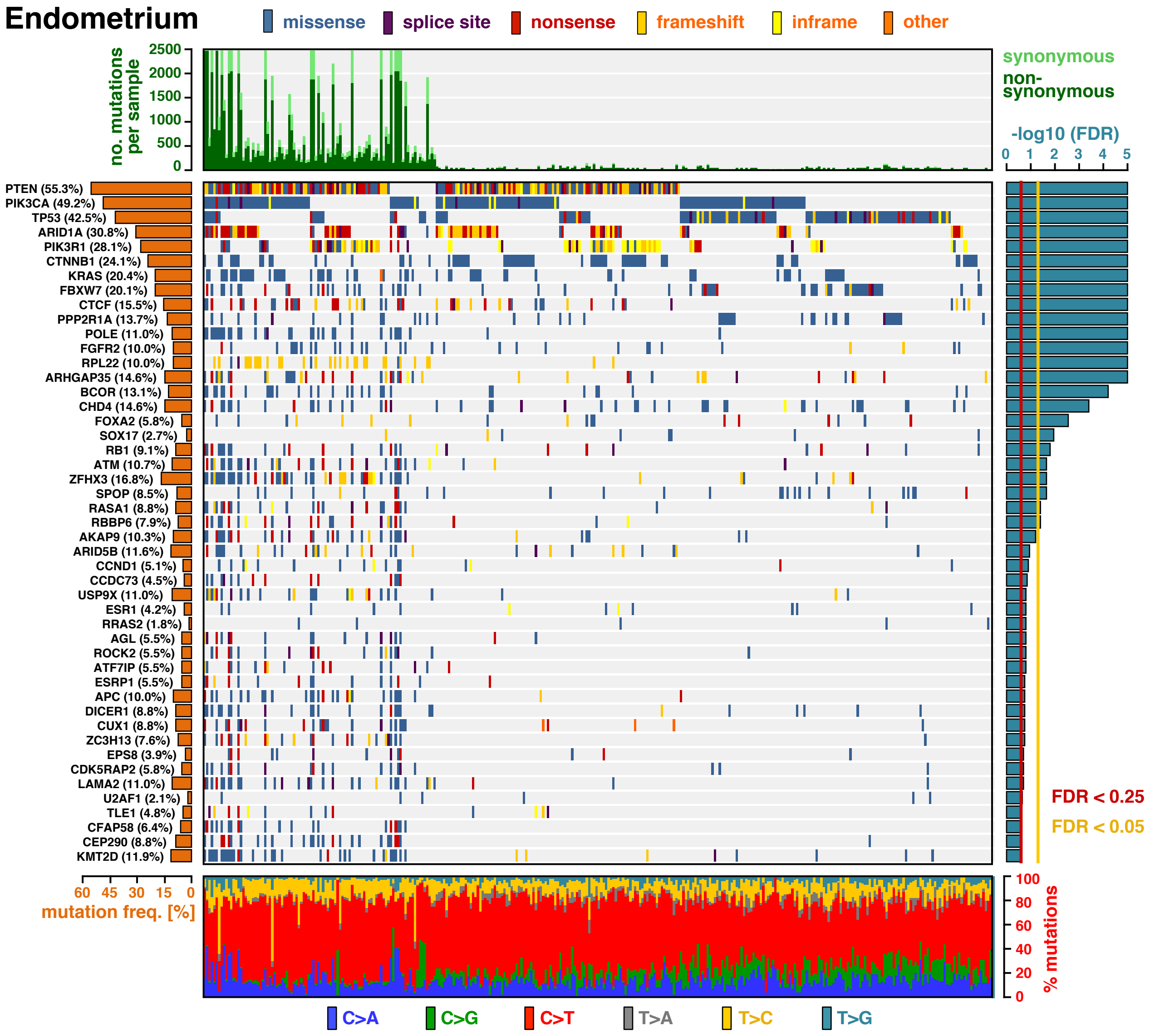


**Extended Data Fig. S26 | The landscape of driver mutations in endometrial carcinomas.** The color-coded matrix in the center displays the distribution of non-synonymous mutations across individual tumor samples (x-axis) in the significantly mutant genes, identified by MutPanning (y-axis). The bar graph on top shows the total number of mutations in each sample. Non-synonymous mutations are colored in dark green; synonymous mutations are colored in light green. The relative contribution of the six mutation types (blue: C>A, green: C>G, red: C>T, gray: T>A, yellow: T>C, cyan: T>G) to the mutational burden of each sample is shown in the bar graph below. The cancer type specific mutation frequency of each significant gene is shown on the left (orange). The False-discovery rates, computed by MutPanning (FDR, q-values), are displayed on the right. The red line indicates a cutoff at q<0.25; the yellow line indicates a threshold at q<0.05.

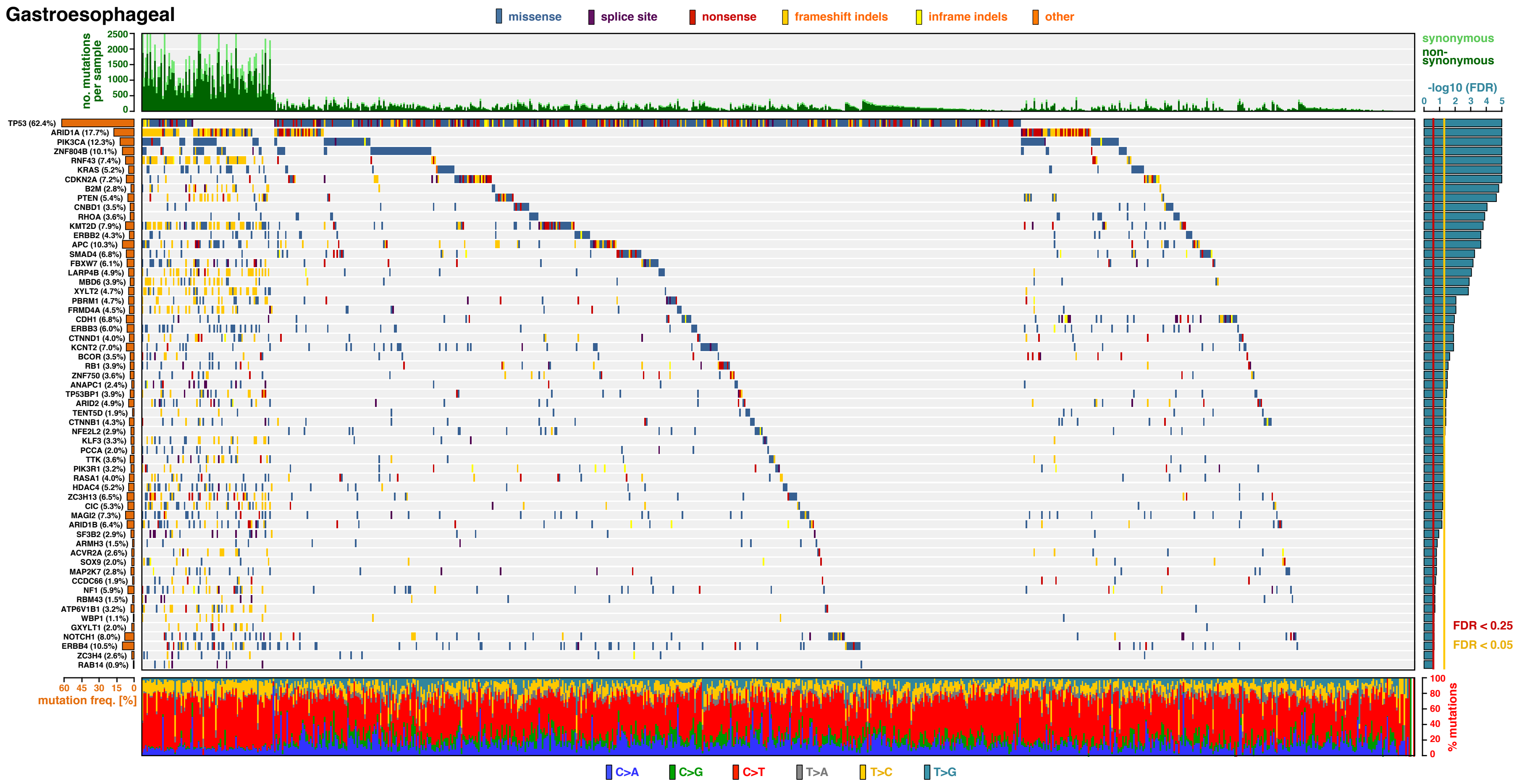


**Extended Data Fig. S27 | The landscape of driver mutations in gastro-esophageal cancer.** The color-coded matrix in the center displays the distribution of non-synonymous mutations across individual tumor samples (x-axis) in the significantly mutant genes, identified by MutPanning (y-axis). The bar graph on top shows the total number of mutations in each sample. Non-synonymous mutations are colored in dark green; synonymous mutations are colored in light green. The relative contribution of the six mutation types (blue: C>A, green: C>G, red: C>T, gray: T>A, yellow: T>C, cyan: T>G) to the mutational burden of each sample is shown in the bar graph below. The cancer type specific mutation frequency of each significant gene is shown on the left (orange). The False-discovery rates, computed by MutPanning (FDR, q-values), are displayed on the right. The red line indicates a cutoff at q<0.25; the yellow line indicates a threshold at q<0.05.

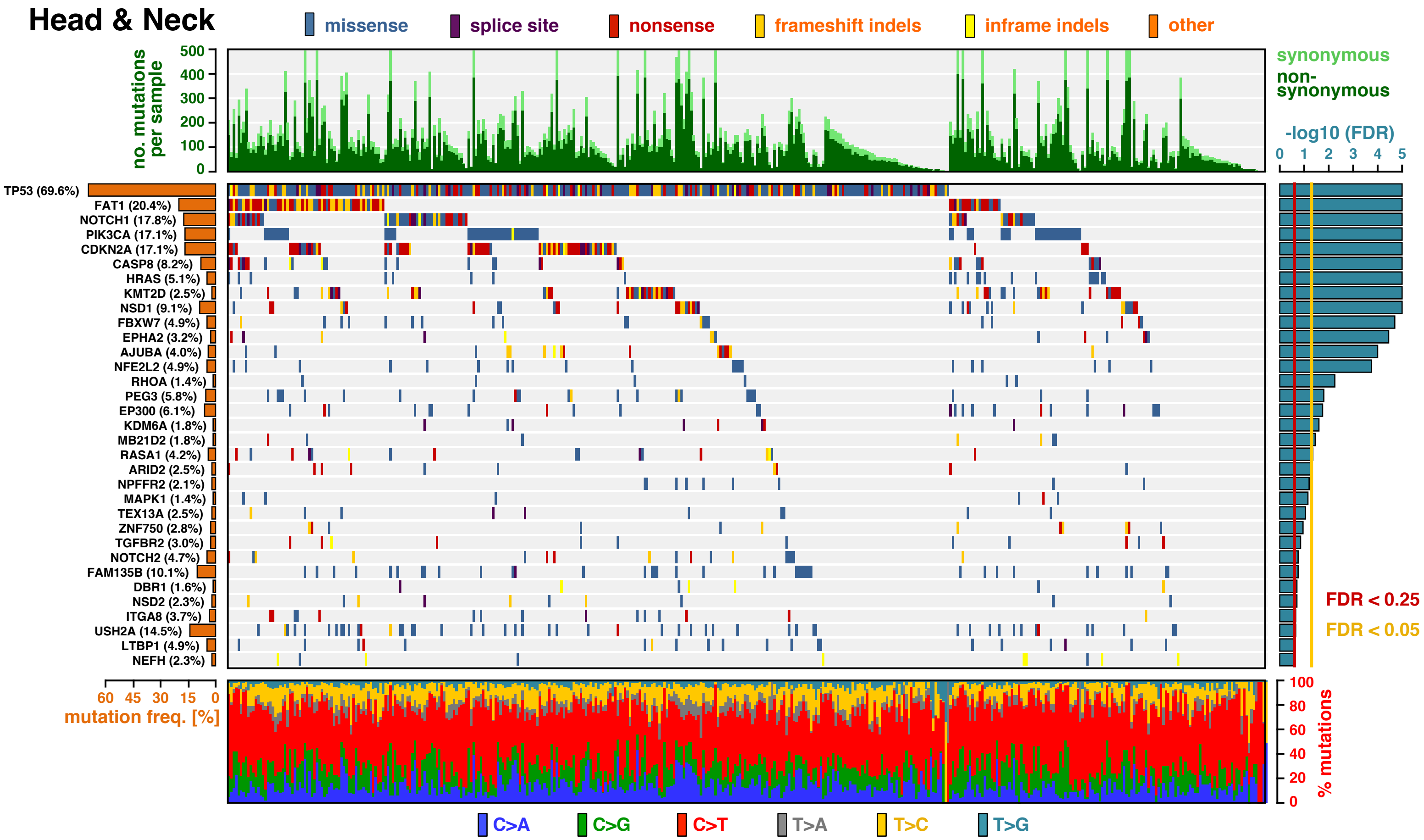


**Extended Data Fig. S28 | The landscape of driver mutations in head and neck carcinoma.** The color-coded matrix in the center displays the distribution of non-synonymous mutations across individual tumor samples (x-axis) in the significantly mutant genes, identified by MutPanning (y-axis). The bar graph on top shows the total number of mutations in each sample. Non-synonymous mutations are colored in dark green; synonymous mutations are colored in light green. The relative contribution of the six mutation types (blue: C>A, green: C>G, red: C>T, gray: T>A, yellow: T>C, cyan: T>G) to the mutational burden of each sample is shown in the bar graph below. The cancer type specific mutation frequency of each significant gene is shown on the left (orange). The False-discovery rates, computed by MutPanning (FDR, q-values), are displayed on the right. The red line indicates a cutoff at q<0.25; the yellow line indicates a threshold at q<0.05.

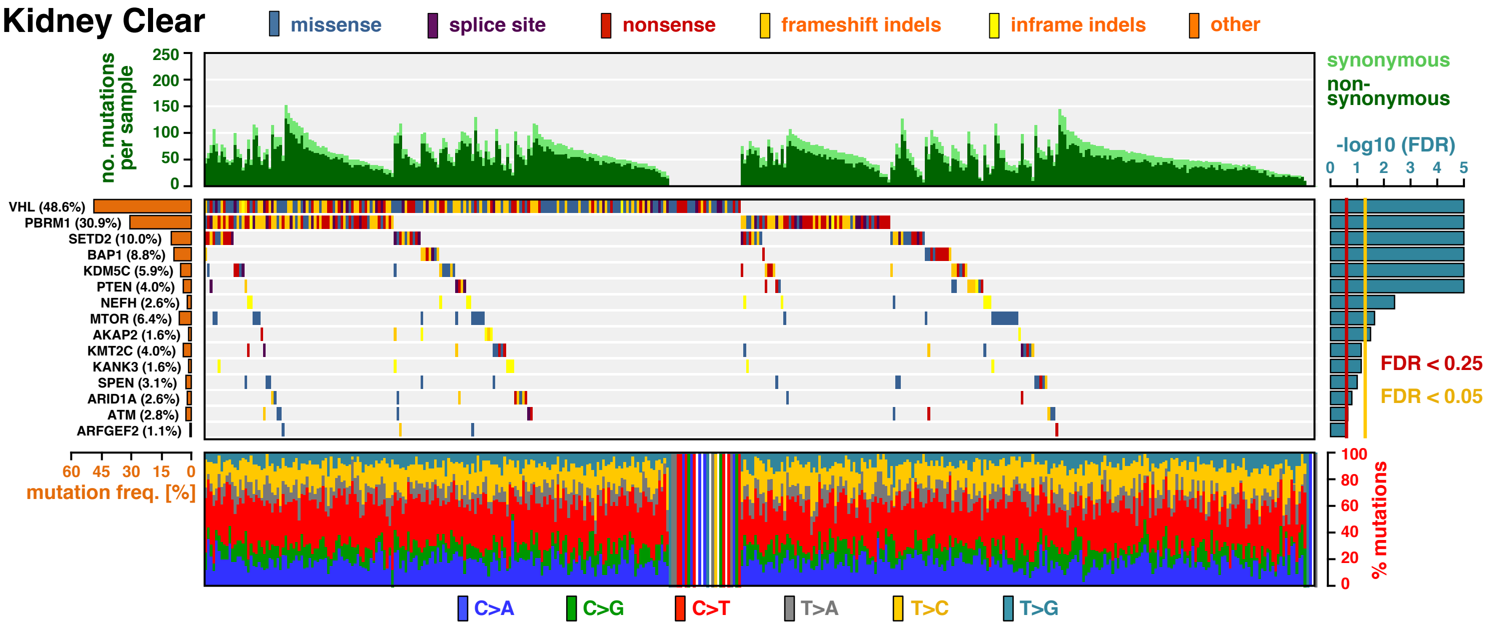


**Extended Data Fig. S29 | The landscape of driver mutations in clear cell kidney cancer.** The color-coded matrix in the center displays the distribution of non-synonymous mutations across individual tumor samples (x-axis) in the significantly mutant genes, identified by MutPanning (y-axis). The bar graph on top shows the total number of mutations in each sample. Non-synonymous mutations are colored in dark green; synonymous mutations are colored in light green. The relative contribution of the six mutation types (blue: C>A, green: C>G, red: C>T, gray: T>A, yellow: T>C, cyan: T>G) to the mutational burden of each sample is shown in the bar graph below. The cancer type specific mutation frequency of each significant gene is shown on the left (orange). The False-discovery rates, computed by MutPanning (FDR, q-values), are displayed on the right. The red line indicates a cutoff at q<0.25; the yellow line indicates a threshold at q<0.05.

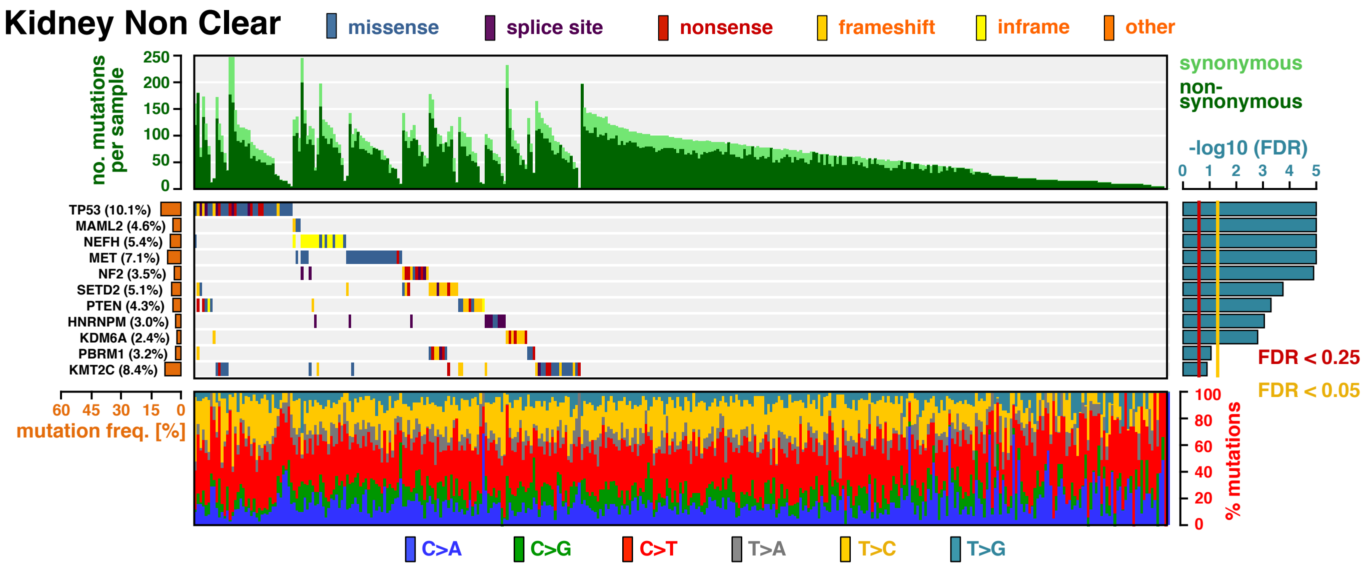


**Extended Data Fig. S30 | The landscape of driver mutations in non-clear cell kidney cancer.** The color-coded matrix in the center displays the distribution of non-synonymous mutations across individual tumor samples (x-axis) in the significantly mutant genes, identified by MutPanning (y-axis). The bar graph on top shows the total number of mutations in each sample. Non-synonymous mutations are colored in dark green; synonymous mutations are colored in light green. The relative contribution of the six mutation types (blue: C>A, green: C>G, red: C>T, gray: T>A, yellow: T>C, cyan: T>G) to the mutational burden of each sample is shown in the bar graph below. The cancer type specific mutation frequency of each significant gene is shown on the left (orange). The False-discovery rates, computed by MutPanning (FDR, q-values), are displayed on the right. The red line indicates a cutoff at q<0.25; the yellow line indicates a threshold at q<0.05.

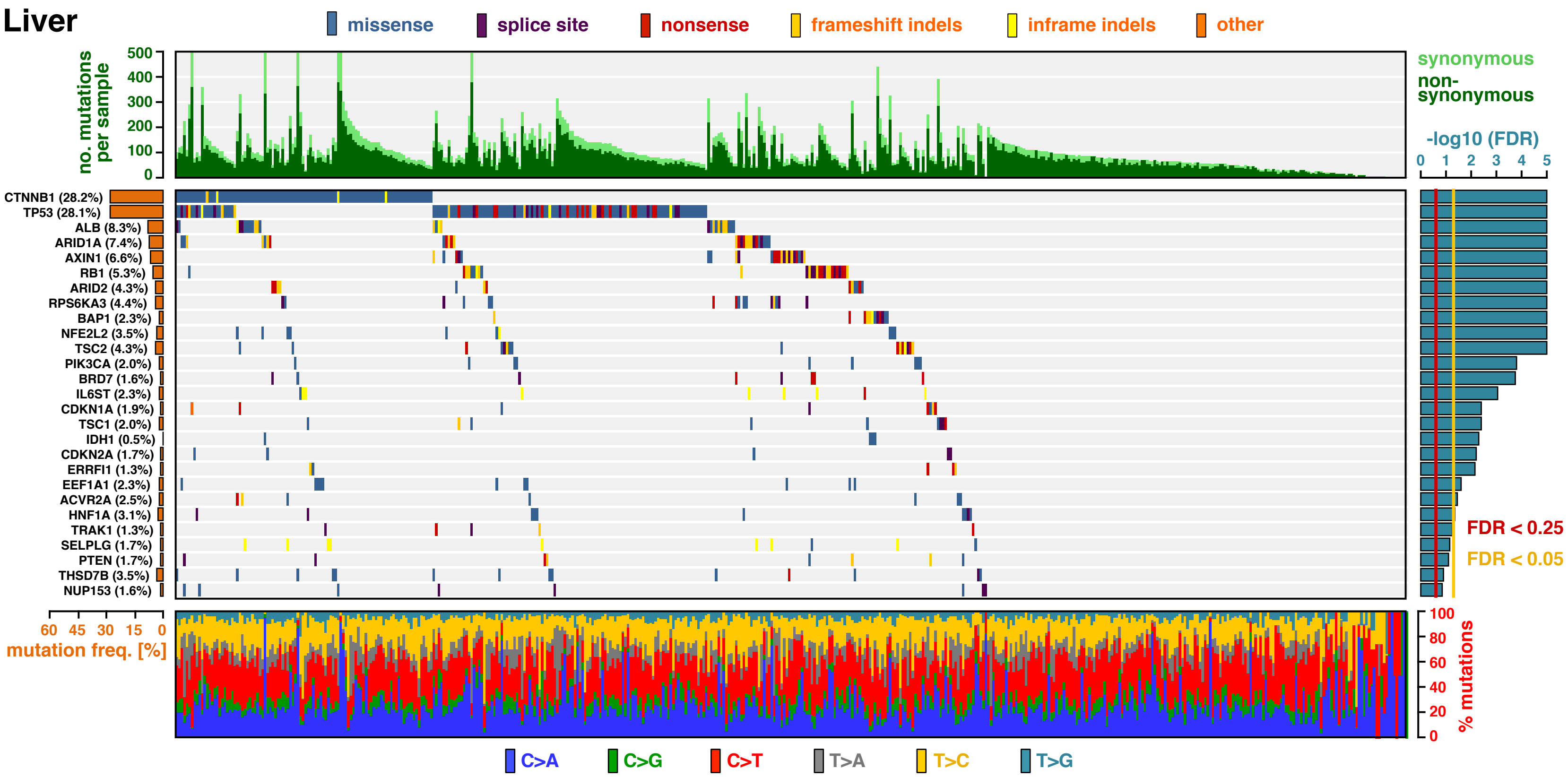


**Extended Data Fig. S31 | The landscape of driver mutations in hepatocellular carcinoma.** The color-coded matrix in the center displays the distribution of non-synonymous mutations across individual tumor samples (x-axis) in the significantly mutant genes, identified by MutPanning (y-axis). The bar graph on top shows the total number of mutations in each sample. Non-synonymous mutations are colored in dark green; synonymous mutations are colored in light green. The relative contribution of the six mutation types (blue: C>A, green: C>G, red: C>T, gray: T>A, yellow: T>C, cyan: T>G) to the mutational burden of each sample is shown in the bar graph below. The cancer type specific mutation frequency of each significant gene is shown on the left (orange). The False-discovery rates, computed by MutPanning (FDR, q-values), are displayed on the right. The red line indicates a cutoff at $q<0.25$; the yellow line indicates a threshold at $q<0.05$.

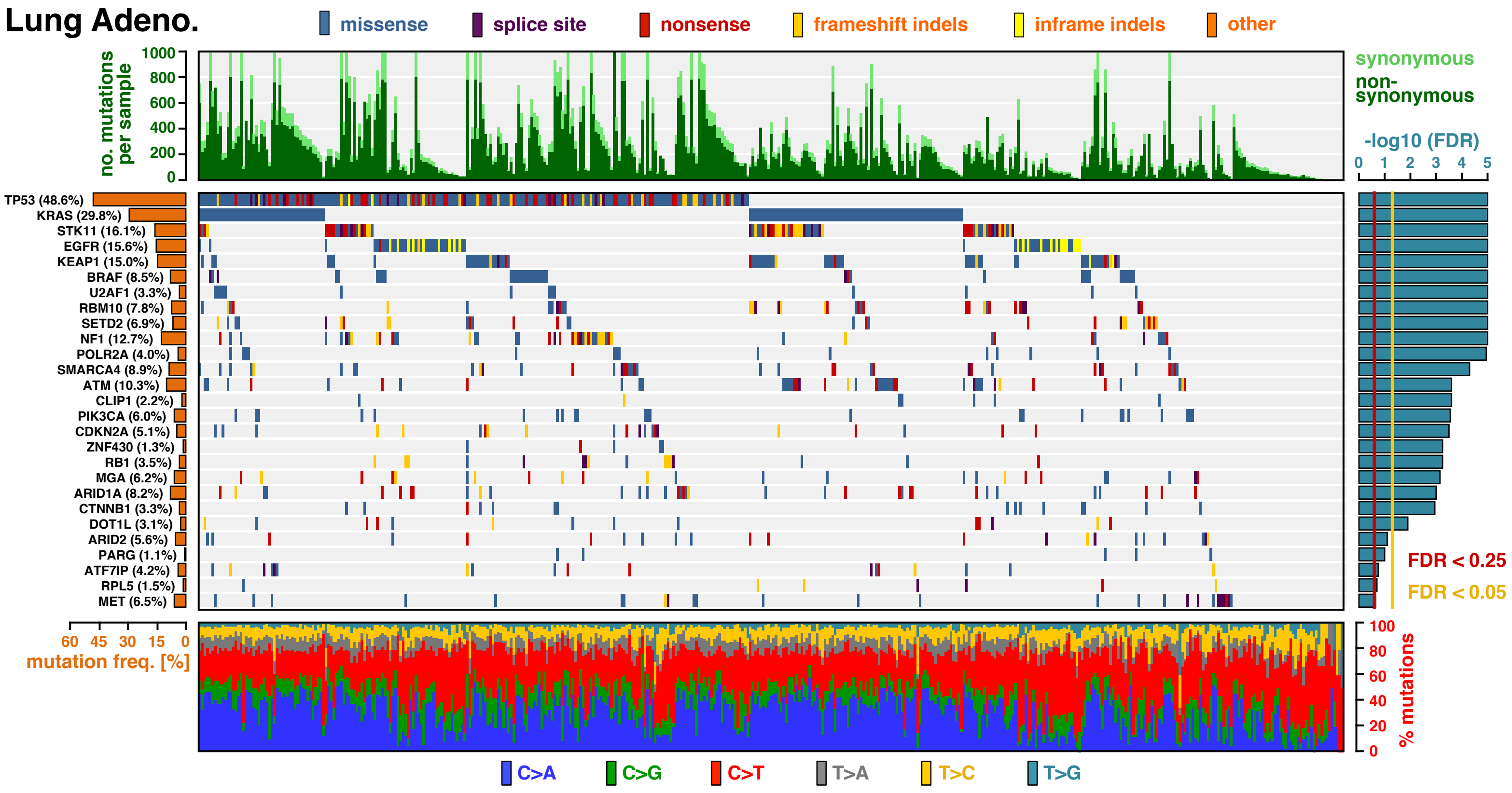


**Extended Data Fig. S32 | The landscape of driver mutations in lung adenocarcinoma.** The color-coded matrix in the center displays the distribution of non-synonymous mutations across individual tumor samples (x-axis) in the significantly mutant genes, identified by MutPanning (y-axis). The bar graph on top shows the total number of mutations in each sample. Non-synonymous mutations are colored in dark green; synonymous mutations are colored in light green. The relative contribution of the six mutation types (blue: C>A, green: C>G, red: C>T, gray: T>A, yellow: T>C, cyan: T>G) to the mutational burden of each sample is shown in the bar graph below. The cancer type specific mutation frequency of each significant gene is shown on the left (orange). The False-discovery rates, computed by MutPanning (FDR, q-values), are displayed on the right. The red line indicates a cutoff at $q<0.25$; the yellow line indicates a threshold at $q<0.05$.

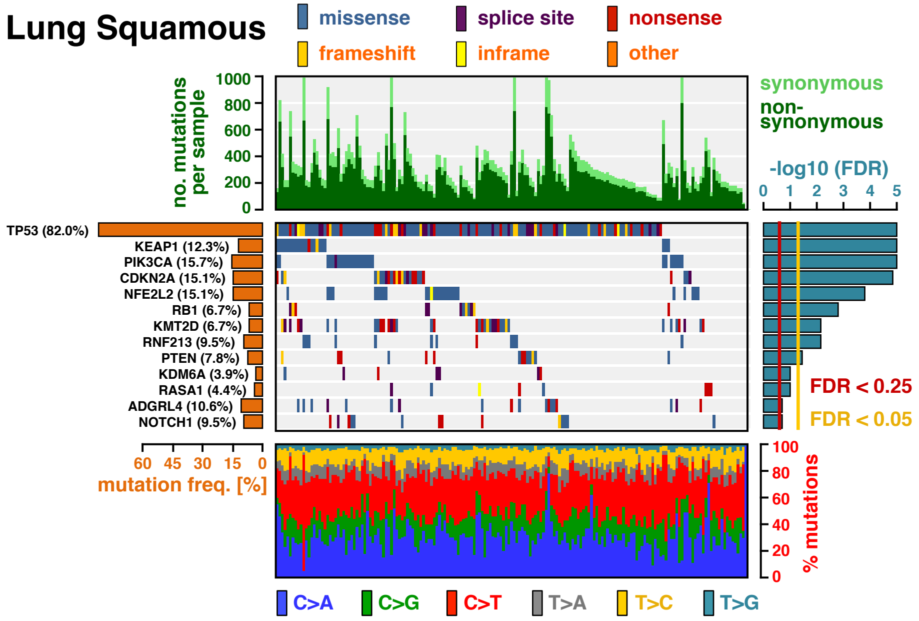


**Extended Data Fig. S33 | The landscape of driver mutations in squamous-cell lung cancer.** The color-coded matrix in the center displays the distribution of non-synonymous mutations across individual tumor samples (x-axis) in the significantly mutant genes, identified by MutPanning (y-axis). The bar graph on top shows the total number of mutations in each sample. Non-synonymous mutations are colored in dark green; synonymous mutations are colored in light green. The relative contribution of the six mutation types (blue: C>A, green: C>G, red: C>T, gray: T>A, yellow: T>C, cyan: T>G) to the mutational burden of each sample is shown in the bar graph below. The cancer type specific mutation frequency of each significant gene is shown on the left (orange). The False-discovery rates, computed by MutPanning (FDR, q-values), are displayed on the right. The red line indicates a cutoff at q<0.25; the yellow line indicates a threshold at q<0.05.

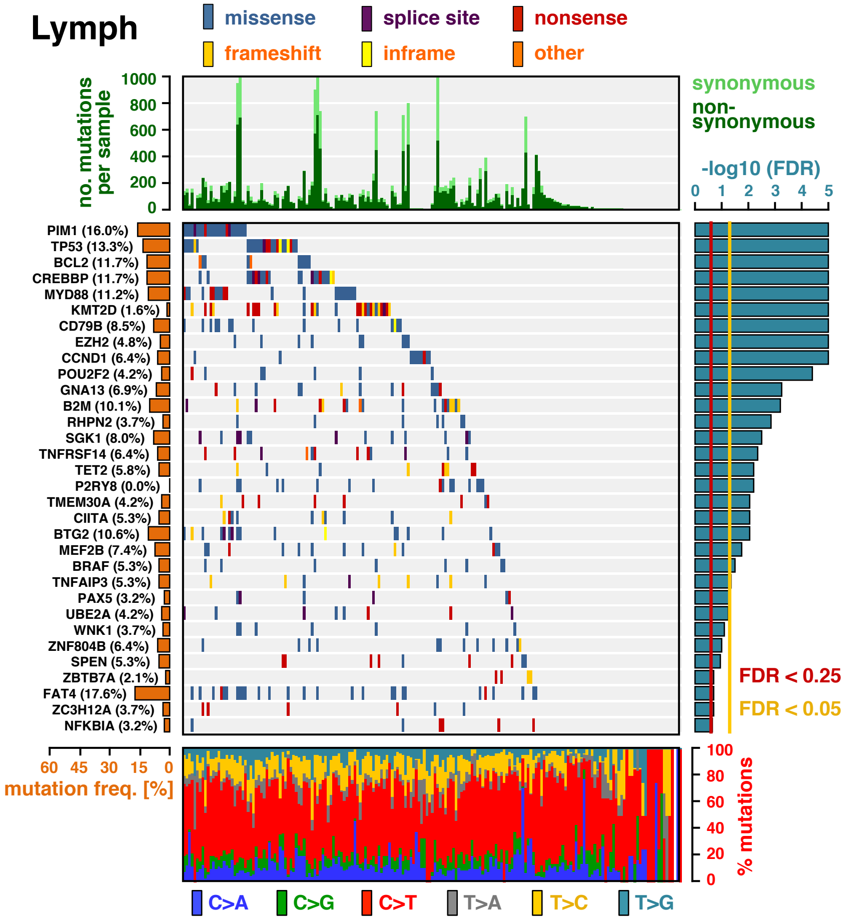


**Extended Data Fig. S34 | The landscape of driver mutations in lymphomas.** The color-coded matrix in the center displays the distribution of non-synonymous mutations across individual tumor samples (x-axis) in the significantly mutant genes, identified by MutPanning (y-axis). The bar graph on top shows the total number of mutations in each sample. Non-synonymous mutations are colored in dark green; synonymous mutations are colored in light green. The relative contribution of the six mutation types (blue: C>A, green: C>G, red: C>T, gray: T>A, yellow: T>C, cyan: T>G) to the mutational burden of each sample is shown in the bar graph below. The cancer type specific mutation frequency of each significant gene is shown on the left (orange). The False-discovery rates, computed by MutPanning (FDR, q-values), are displayed on the right. The red line indicates a cutoff at q<0.25; the yellow line indicates a threshold at q<0.05.

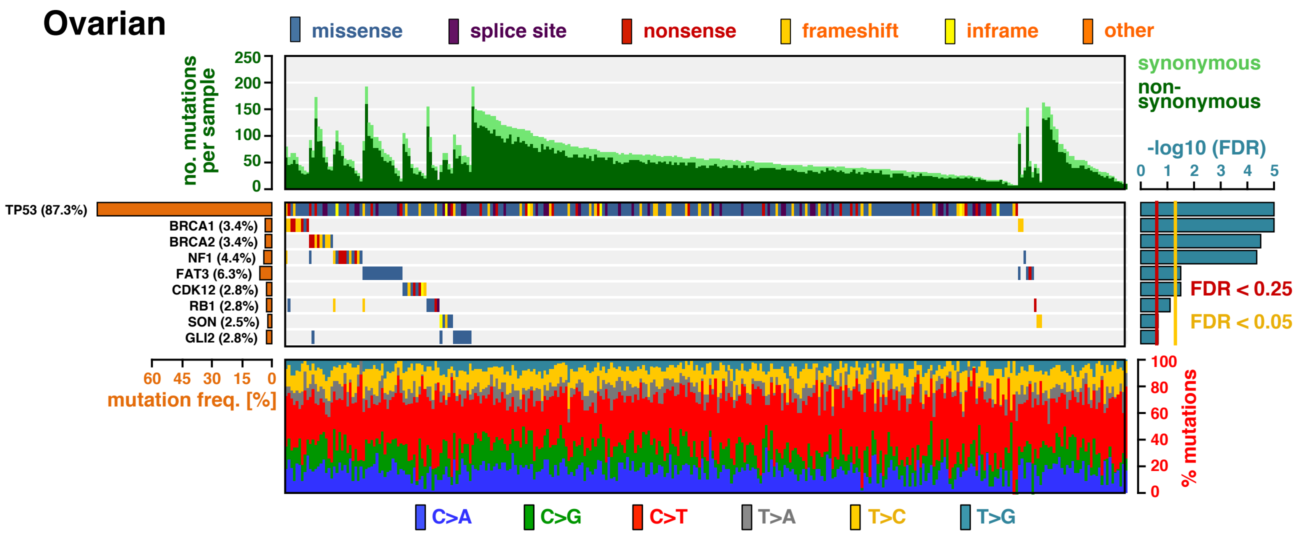


**Extended Data Fig. S35 | The landscape of driver mutations in ovarian cancer.** The color-coded matrix in the center displays the distribution of non-synonymous mutations across individual tumor samples (x-axis) in the significantly mutant genes, identified by MutPanning (y-axis). The bar graph on top shows the total number of mutations in each sample. Non-synonymous mutations are colored in dark green; synonymous mutations are colored in light green. The relative contribution of the six mutation types (blue: C>A, green: C>G, red: C>T, gray: T>A, yellow: T>C, cyan: T>G) to the mutational burden of each sample is shown in the bar graph below. The cancer type specific mutation frequency of each significant gene is shown on the left (orange). The False-discovery rates, computed by MutPanning (FDR, q-values), are displayed on the right. The red line indicates a cutoff at q<0.25; the yellow line indicates a threshold at q<0.05.

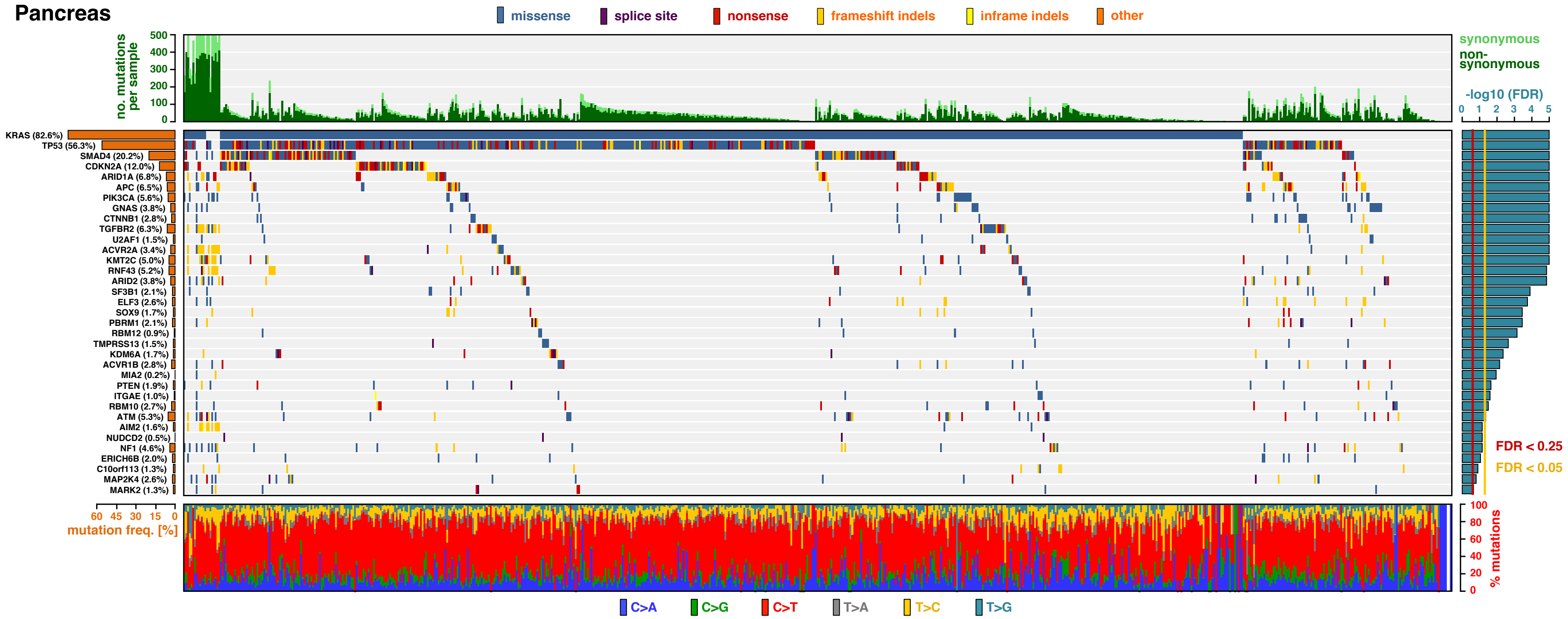


**Extended Data Fig. S36 | The landscape of driver mutations in pancreatic cancer.** The color-coded matrix in the center displays the distribution of non-synonymous mutations across individual tumor samples (x-axis) in the significantly mutant genes, identified by MutPanning (y-axis). The bar graph on top shows the total number of mutations in each sample. Non-synonymous mutations are colored in dark green; synonymous mutations are colored in light green. The relative contribution of the six mutation types (blue: C>A, green: C>G, red: C>T, gray: T>A, yellow: T>C, cyan: T>G) to the mutational burden of each sample is shown in the bar graph below. The cancer type specific mutation frequency of each significant gene is shown on the left (orange). The False-discovery rates, computed by MutPanning (FDR, q-values), are displayed on the right. The red line indicates a cutoff at q<0.25; the yellow line indicates a threshold at q<0.05.

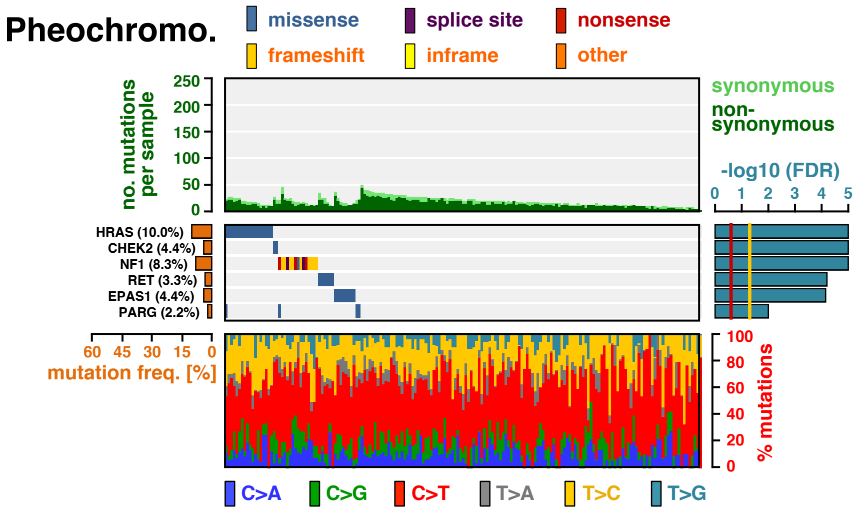


**Extended Data Fig. S37 | The landscape of driver mutations in pheochromocytomas and paragangliomas.** The color-coded matrix in the center displays the distribution of non-synonymous mutations across individual tumor samples (x-axis) in the significantly mutant genes, identified by MutPanning (y-axis). The bar graph on top shows the total number of mutations in each sample. Non-synonymous mutations are colored in dark green; synonymous mutations are colored in light green. The relative contribution of the six mutation types (blue: C>A, green: C>G, red: C>T, gray: T>A, yellow: T>C, cyan: T>G) to the mutational burden of each sample is shown in the bar graph below. The cancer type specific mutation frequency of each significant gene is shown on the left (orange). The False-discovery rates, computed by MutPanning (FDR, q-values), are displayed on the right. The red line indicates a cutoff at q<0.25; the yellow line indicates a threshold at q<0.05.

### Pleura

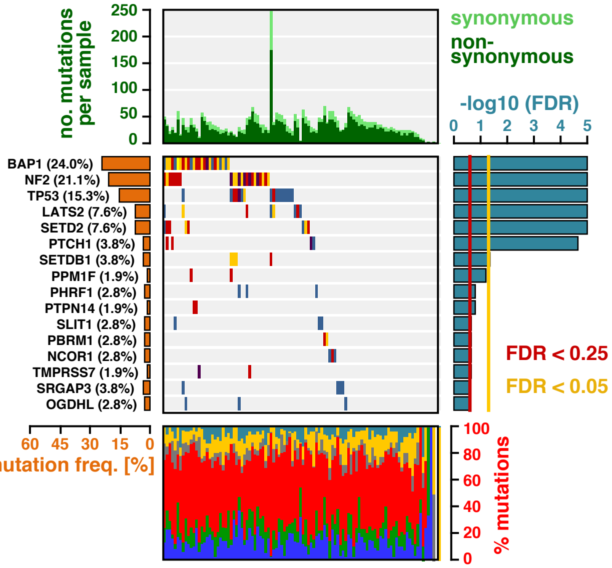


**Extended Data Fig. S38 | The landscape of driver mutations in mesothelioma.** The color-coded matrix in the center displays the distribution of non-synonymous mutations across individual tumor samples (x-axis) in the significantly mutant genes, identified by MutPanning (y-axis). The bar graph on top shows the total number of mutations in each sample. Non-synonymous mutations are colored in dark green; synonymous mutations are colored in light green. The relative contribution of the six mutation types (blue: C>A, green: C>G, red: C>T, gray: T>A, yellow: T>C, cyan: T>G) to the mutational burden of each sample is shown in the bar graph below. The cancer type specific mutation frequency of each significant gene is shown on the left (orange). The False-discovery rates, computed by MutPanning (FDR, q-values), are displayed on the right. The red line indicates a cutoff at q<0.25; the yellow line indicates a threshold at q<0.05.

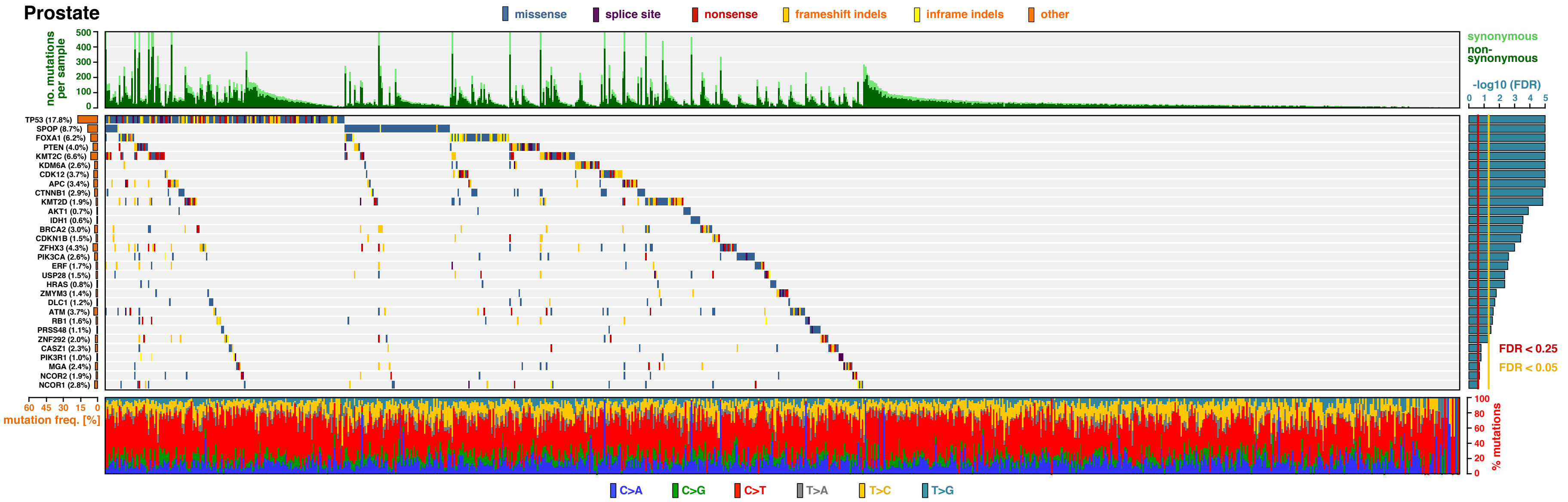


**Extended Data Fig. S39 | The landscape of driver mutations in prostate cancer.** The color-coded matrix in the center displays the distribution of non-synonymous mutations across individual tumor samples (x-axis) in the significantly mutant genes, identified by MutPanning (y-axis). The bar graph on top shows the total number of mutations in each sample. Non-synonymous mutations are colored in dark green; synonymous mutations are colored in light green. The relative contribution of the six mutation types (blue: C>A, green: C>G, red: C>T, gray: T>A, yellow: T>C, cyan: T>G) to the mutational burden of each sample is shown in the bar graph below. The cancer type specific mutation frequency of each significant gene is shown on the left (orange). The False-discovery rates, computed by MutPanning (FDR, q-values), are displayed on the right. The red line indicates a cutoff at q<0.25; the yellow line indicates a threshold at q<0.05.

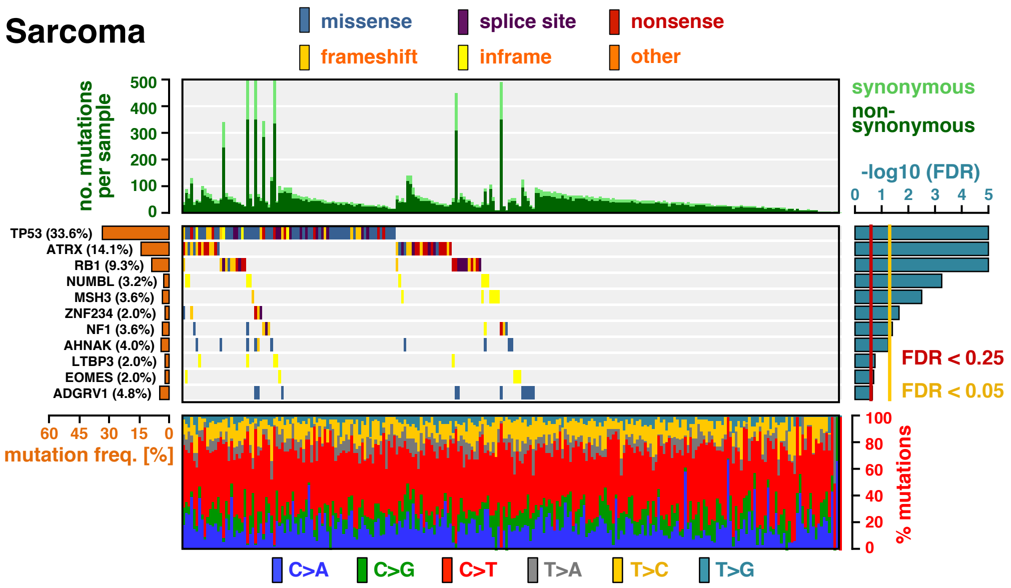


**Extended Data Fig. S40 | The landscape of driver mutations in adult soft tissue sarcoma.** The color-coded matrix in the center displays the distribution of non-synonymous mutations across individual tumor samples (x-axis) in the significantly mutant genes, identified by MutPanning (y-axis). The bar graph on top shows the total number of mutations in each sample. Non-synonymous mutations are colored in dark green; synonymous mutations are colored in light green. The relative contribution of the six mutation types (blue: C>A, green: C>G, red: C>T, gray: T>A, yellow: T>C, cyan: T>G) to the mutational burden of each sample is shown in the bar graph below. The cancer type specific mutation frequency of each significant gene is shown on the left (orange). The False-discovery rates, computed by MutPanning (FDR, q-values), are displayed on the right. The red line indicates a cutoff at q<0.25; the yellow line indicates a threshold at q<0.05.

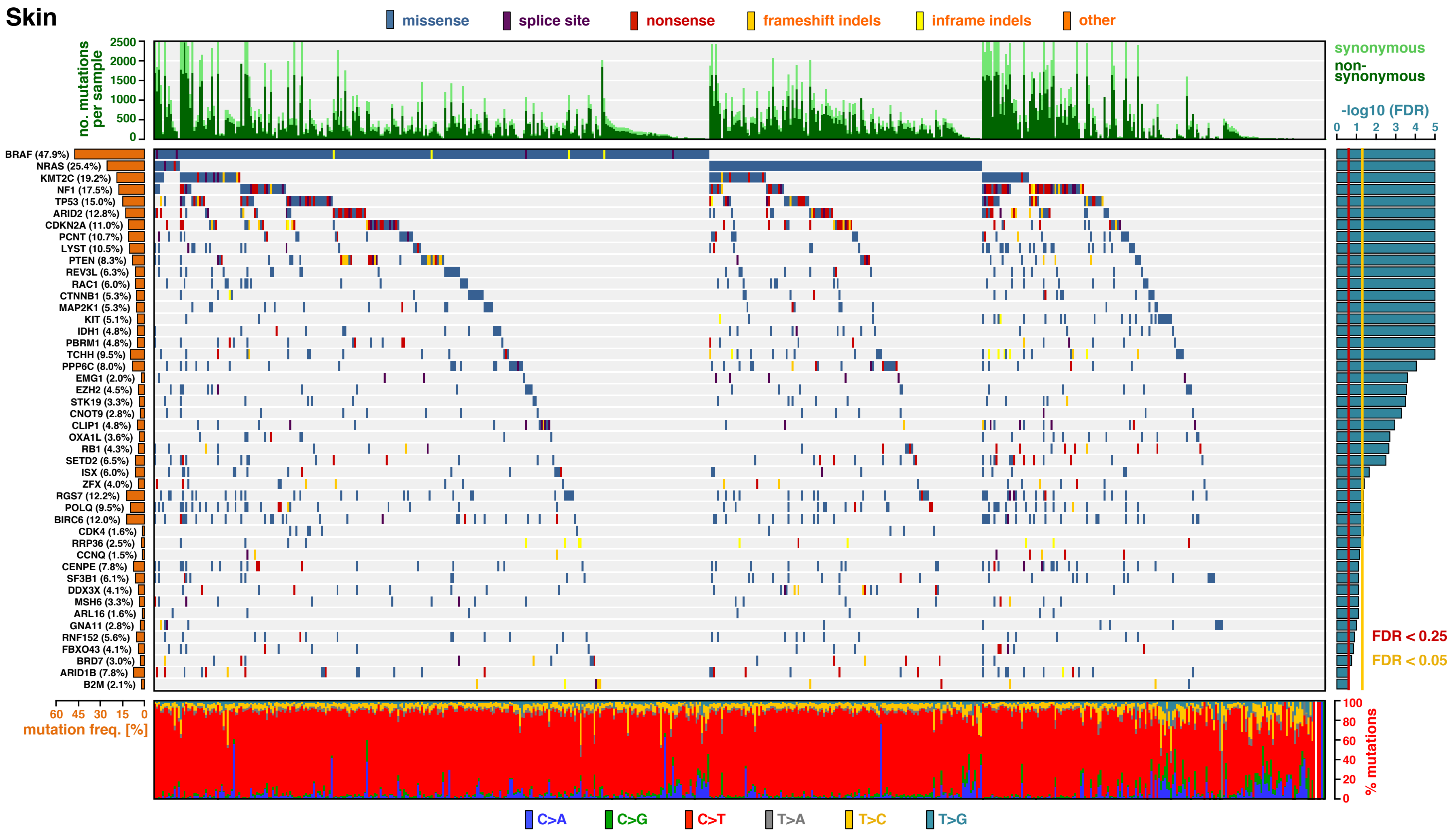


**Extended Data Fig. S41 | The landscape of driver mutations in melanoma.** The color-coded matrix in the center displays the distribution of non-synonymous mutations across individual tumor samples (x-axis) in the significantly mutant genes, identified by MutPanning (y-axis). The bar graph on top shows the total number of mutations in each sample. Non-synonymous mutations are colored in dark green; synonymous mutations are colored in light green. The relative contribution of the six mutation types (blue: C>A, green: C>G, red: C>T, gray: T>A, yellow: T>C, cyan: T>G) to the mutational burden of each sample is shown in the bar graph below. The cancer type specific mutation frequency of each significant gene is shown on the left (orange). The False-discovery rates, computed by MutPanning (FDR, q-values), are displayed on the right. The red line indicates a cutoff at q<0.25; the yellow line indicates a threshold at q<0

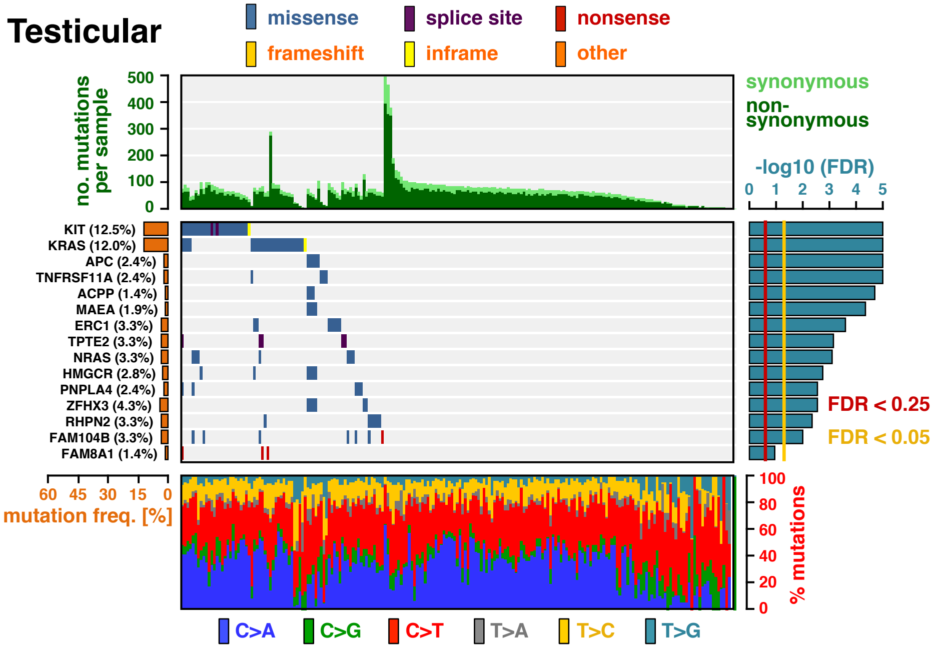


**Extended Data Fig. S42 | The landscape of driver mutations in testicular germ cell tumors.** The color-coded matrix in the center displays the distribution of non-synonymous mutations across individual tumor samples (x-axis) in the significantly mutant genes, identified by MutPanning (y-axis). The bar graph on top shows the total number of mutations in each sample. Non-synonymous mutations are colored in dark green; synonymous mutations are colored in light green. The relative contribution of the six mutation types (blue: C>A, green: C>G, red: C>T, gray: T>A, yellow: T>C, cyan: T>G) to the mutational burden of each sample is shown in the bar graph below. The cancer type specific mutation frequency of each significant gene is shown on the left (orange). The False-discovery rates, computed by MutPanning (FDR, q-values), are displayed on the right. The red line indicates a cutoff at q<0.25; the yellow line indicates a threshold at q<0.05.

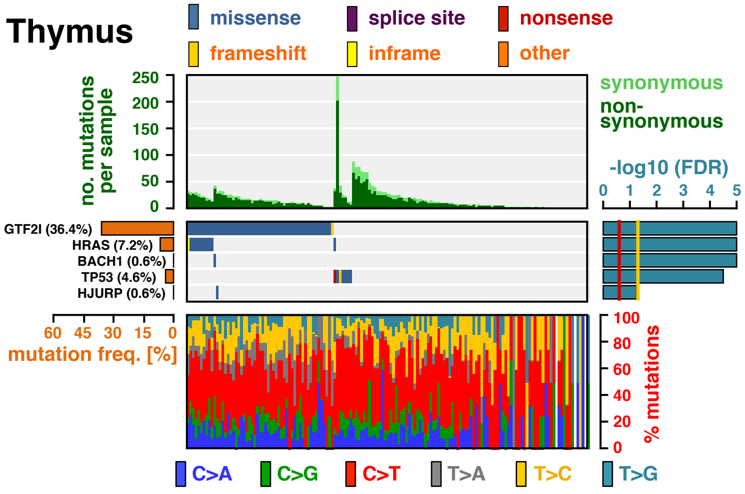


**Extended Data Fig. S43 | The landscape of driver mutations in thymic epithelial tumors.** The color-coded matrix in the center displays the distribution of non-synonymous mutations across individual tumor samples (x-axis) in the significantly mutant genes, identified by MutPanning (y-axis). The bar graph on top shows the total number of mutations in each sample. Non-synonymous mutations are colored in dark green; synonymous mutations are colored in light green. The relative contribution of the six mutation types (blue: C>A, green: C>G, red: C>T, gray: T>A, yellow: T>C, cyan: T>G) to the mutational burden of each sample is shown in the bar graph below. The cancer type specific mutation frequency of each significant gene is shown on the left (orange). The False-discovery rates, computed by MutPanning (FDR, q-values), are displayed on the right. The red line indicates a cutoff at q<0.25; the yellow line indicates a threshold at q<0.05.

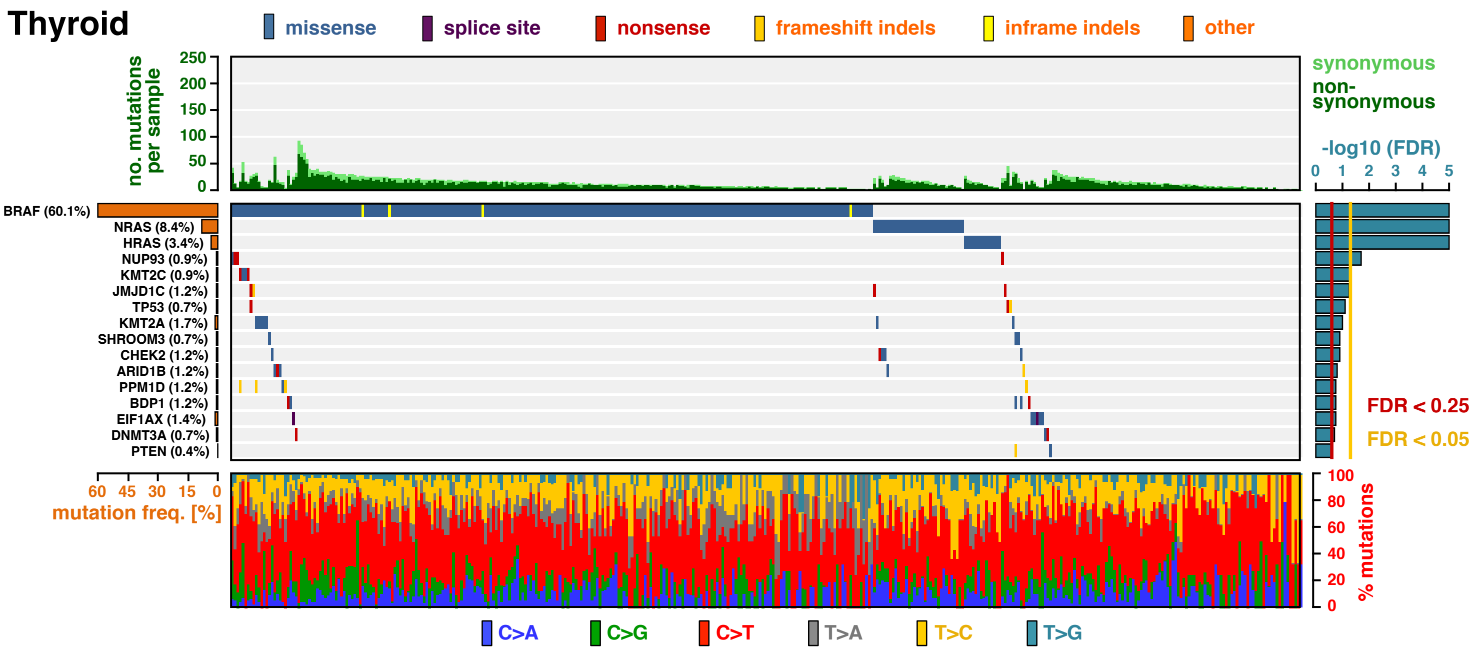


**Extended Data Fig. S44 | The landscape of driver mutations in thyroid cancer.** The color-coded matrix in the center displays the distribution of non-synonymous mutations across individual tumor samples (x-axis) in the significantly mutant genes, identified by MutPanning (y-axis). The bar graph on top shows the total number of mutations in each sample. Non-synonymous mutations are colored in dark green; synonymous mutations are colored in light green. The relative contribution of the six mutation types (blue: C>A, green: C>G, red: C>T, gray: T>A, yellow: T>C, cyan: T>G) to the mutational burden of each sample is shown in the bar graph below. The cancer type specific mutation frequency of each significant gene is shown on the left (orange). The False-discovery rates, computed by MutPanning (FDR, q-values), are displayed on the right. The red line indicates a cutoff at q<0.25; the yellow line indicates a threshold at q<0.05.

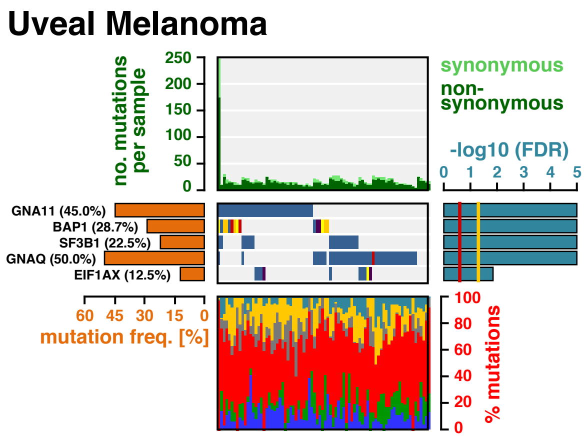


**Extended Data Fig. S45 | The landscape of driver mutations in uveal melanoma.** The color-coded matrix in the center displays the distribution of non-synonymous mutations across individual tumor samples (x-axis) in the significantly mutant genes, identified by MutPanning (y-axis). The bar graph on top shows the total number of mutations in each sample. Non-synonymous mutations are colored in dark green; synonymous mutations are colored in light green. The relative contribution of the six mutation types (blue: C>A, green: C>G, red: C>T, gray: T>A, yellow: T>C, cyan: T>G) to the mutational burden of each sample is shown in the bar graph below. The cancer type specific mutation frequency of each significant gene is shown on the left (orange). The False-discovery rates, computed by MutPanning (FDR, q-values), are displayed on the right. The red line indicates a cutoff at q<0.25; the yellow line indicates a threshold at q<0.05.
